# reconcILS: A gene tree-species tree reconciliation algorithm that allows for incomplete lineage sorting

**DOI:** 10.1101/2023.11.03.565544

**Authors:** Sarthak R. Mishra, Megan L. Smith, Matthew W. Hahn

## Abstract

Reconciliation algorithms infer the evolutionary history of individual gene trees given a species tree. Many reconciliation algorithms consider only duplication and loss events (and sometimes horizontal transfer), ignoring effects of the coalescent process, including incomplete lineage sorting (ILS). Here, we present a new parsimony algorithm for carrying out reconciliation that accurately accounts for ILS by treating it as a series of nearest neighbor interchange (NNI) events on a tree. For discordant branches of the gene tree, our algorithm recursively chooses the optimal history by comparing the cost of duplication and loss to the cost of NNI and loss. We demonstrate the accuracy of our new method, which we call `reconcILS`, using a new simulation engine (`dupcoal`) that generates gene trees produced by the interaction of duplication, loss, and coalescence under the MSC-DL model. Despite being a heuristic method, `reconcILS` is much more accurate than models that ignore ILS, and at least as accurate or better than leading methods that can model ILS, while also being able to handle larger datasets. We demonstrate the use of `reconcILS` by applying it to a dataset of 23 primate genomes, highlighting its accuracy compared to standard methods in the presence of large amounts of ILS.

## Introduction

The sequencing of a large number of genomes from many different species has highlighted the dynamic nature of genes: they are often duplicated, lost, or moved between species via introgression or horizontal gene transfer. One of the main goals of evolutionary and comparative genomics has been to quantify this dynamism, both in terms of the number of events of each type and their timing. A popular approach for quantifying both measures is gene tree-species tree reconciliation (reviewed in Boussau and Scornavacca 2020). Given a gene tree topology and a species tree topology, reconciliation algorithms attempt to infer the evolutionary events necessary to explain any discordance in the respective topologies. Many different reconciliation algorithms have been developed, most often considering only duplication and loss (so-called “DL” models; e.g. Goodman et al. 1979; Page 1994; Guigo et al. 1996; Page and Charleston 1997; Chen et al. 2000; Zmasek and Eddy 2001; Durand et al. 2006), but also sometimes considering duplication, transfer, and loss (“DTL” models; e.g. Hallett and Lagergren 2001; Hallett et al. 2004; Bansal et al. 2012; Stolzer et al. 2012; Szöllősi et al. 2013; Jacox et al. 2016; Morel et al. 2020).

Despite the success—and widespread use—of DL and DTL reconciliation algorithms, they do not consider gene tree discordance due to the coalescent process. The coalescent is a model that describes how sequences within a population are related (Kingman 1982; Hudson 1983; Tajima 1983), and is relevant even when considering sequences from different species because these must also coalesce in shared ancestral populations. Importantly, gene trees and species trees can be discordant solely due to coalescence, in a process often called incomplete lineage sorting (ILS) or deep coalescence (Maddison 1997). Standard DL and DTL algorithms will incorrectly infer one duplication and three losses for every discordance due only to ILS (Hahn 2007), resulting in misleading results when applying reconciliation methods to trees with ILS (e.g. Stull et al. 2021). Duplication, loss, and coalescence can also interact in complex and non-intuitive ways (Rasmussen and Kellis 2012; Mallo et al. 2014; Li et al. 2021; Mishra and Hahn 2026), making it almost impossible to simply exclude gene trees discordant due to ILS from an analysis.

A few different approaches to reconciliation have been used to accommodate discordance due to coalescence. An early approach allowed short branches in the species tree—which can lead to ILS—to be collapsed (Vernot et al. 2008; Stolzer et al. 2012). The use of non-binary species trees means that such algorithms will not mistake ILS alone for duplication and loss; however, they do not account for all effects of coalescence on incorrect inferences of duplication and loss (see below). Chan et al. (2017) introduced a reconciliation algorithm that included duplication, loss, transfer, and ILS, but it assumed that new duplicates were completely linked to their parental locus (i.e. no recombination between them). Given that the majority of new gene duplicates are not completely linked (Schrider and Hahn 2010), such an assumption limits the number of genes that can be analyzed. Rasmussen and Kellis (2012) introduced a probabilistic model considering duplication, loss, and ILS and an associated reconciliation algorithm. While promising, DLCoalRecon is highly parameterized, requiring users to specify rates of duplication and loss, population sizes, and more prior to inference, and is computationally intensive. Wu et al. (2014) developed a parsimony-based algorithm (DLCpar) based on this same model that runs faster and without the need to specify duplication and loss rates ahead of time.

Here, we introduce a new heuristic method for finding the most parsimonious reconciliation of a gene tree and a species tree in the presence of duplication, loss, and ILS. Our goal is to improve upon existing methods by increasing speed and accuracy, as well as by incorporating more coalescent processes (i.e. ILS is not the only cause of discordance due to the coalescent—see section below on “Complex coalescence”). In addition, although DLCpar has been shown to perform well on smaller trees (Wu et al. 2014; Du et al. 2021), it is not straightforward to run it on larger datasets or to include its algorithm within other tools that also have reconciliation steps (e.g. Thomas et al. 2017; Zhang et al. 2020). We hope that the algorithm introduced here will be easily adaptable to such scenarios. We demonstrate the use of this new algorithm by applying it to 23 whole genomes from primates, a dataset that has previously been shown to contain a large amount of discordance due to ILS (Vanderpool et al. 2020).

## New Approaches

### The multispecies coalescent model with duplication and loss

We begin with a description of the multispecies coalescent model with duplication and loss (MSC-DL). Our model is very similar to the multilocus multispecies coalescent (MLMSC) model of Li et al. (2021). The description of the MLMSC model and an algorithm for generating trees under this model were first given in Li et al. (2021), with a mathematical model and further algorithmic updates in Li et al. (2024). The Appendix (section 1) goes over the conceptual and algorithmic differences between the MSC-DL and MLMSC. Below, we describe the most important biological parts of our model. This explanation is intended to make clearer which common biological processes are modeled by the MSC-DL, but not by standard DL and DTL methods (see also Mishra and Hahn 2026).

The MSC-DL model treats each individual locus (i.e. non-recombing position in the genome) as evolving via the multispecies coalescent (MSC) model, in which discordance can arise only via incomplete lineage sorting. The MSC-DL further models relationships among these loci via duplication and coalescence, in addition to losses at any locus (Figure 1; Supplementary Figure S1). In particular, the MSC-DL specifies:

i. Every locus in the genome has a history that follows the species tree. This specification only becomes important if introgression is allowed (i.e. under the parent-tree model of Meng and Kubatko 2009), but we state it here for completeness.
ii. Each locus in the genome has a gene tree topology drawn from the species tree according to the MSC model. This gene tree (or “genealogy”) exists at this locus whether or not a duplication or loss has occurred; i.e. it is equivalent to the gene tree at any single-copy locus.
iii. Duplication events copy genetic material—a DNA sequence—from one locus (the “parent” locus) and insert it at another locus (the “daughter” locus). At the parent locus, duplicated genetic material descends from a lineage that existed at the time of duplication (star in Figure 1a; filled circle in Supplementary Figure S2a), and so must coalesce above this point with other lineages at the parent locus according to the MSC. At the daughter locus, this material inserts into (is inherited by) a randomly chosen lineage of the gene tree at the time of duplication (e.g. lineage *C* in Figure 1b), following the gene tree topology of this lineage after this point. Nucleotide mutations occur in the sequence that was duplicated, allowing us to reconstruct gene trees connecting one or more loci.
iv. Parent and daughter loci can be any recombination distance apart. If they are completely linked they share the same gene tree topology, while if they are unlinked each locus has an independent topology.
v. Duplications and losses are generated at rates λ and μ via a birth-death process, starting at the original locus. Each duplication event adds a sub-tree from a new daughter locus, consisting only of lineages descending from the duplication (see point iii). Losses remove descendant branches of gene trees below their occurrence (see point vii).
vi. Duplications and losses can occur on any gene tree lineage at any locus that contains genetic material descended from the original parent locus, regardless of whether they were the original parent or daughter locus.
vii. Loss events do not remove a locus, but rather do not allow lineages descended from a loss to extend to the present (Supplementary Figure S3). These lineages are therefore not present in the full gene tree.
viii. The gene tree at each daughter locus coalesces with the gene tree at its parent locus via the MSC (Figure 1c-g; Supplementary Figures S1, S2). This coalescent event occurs further in the past than the duplication event, as the duplicated sequence already has a history with other lineages at the parent locus. Note that even if parent and daughter loci are completely linked, they still must coalesce together at the parent locus—linkage only determines whether the two marginal gene trees have the same topology or not.
ix. The “full” gene tree consists of the combined genealogies at all parent and daughter loci together, including all duplication and loss events.

**Figure 1.**
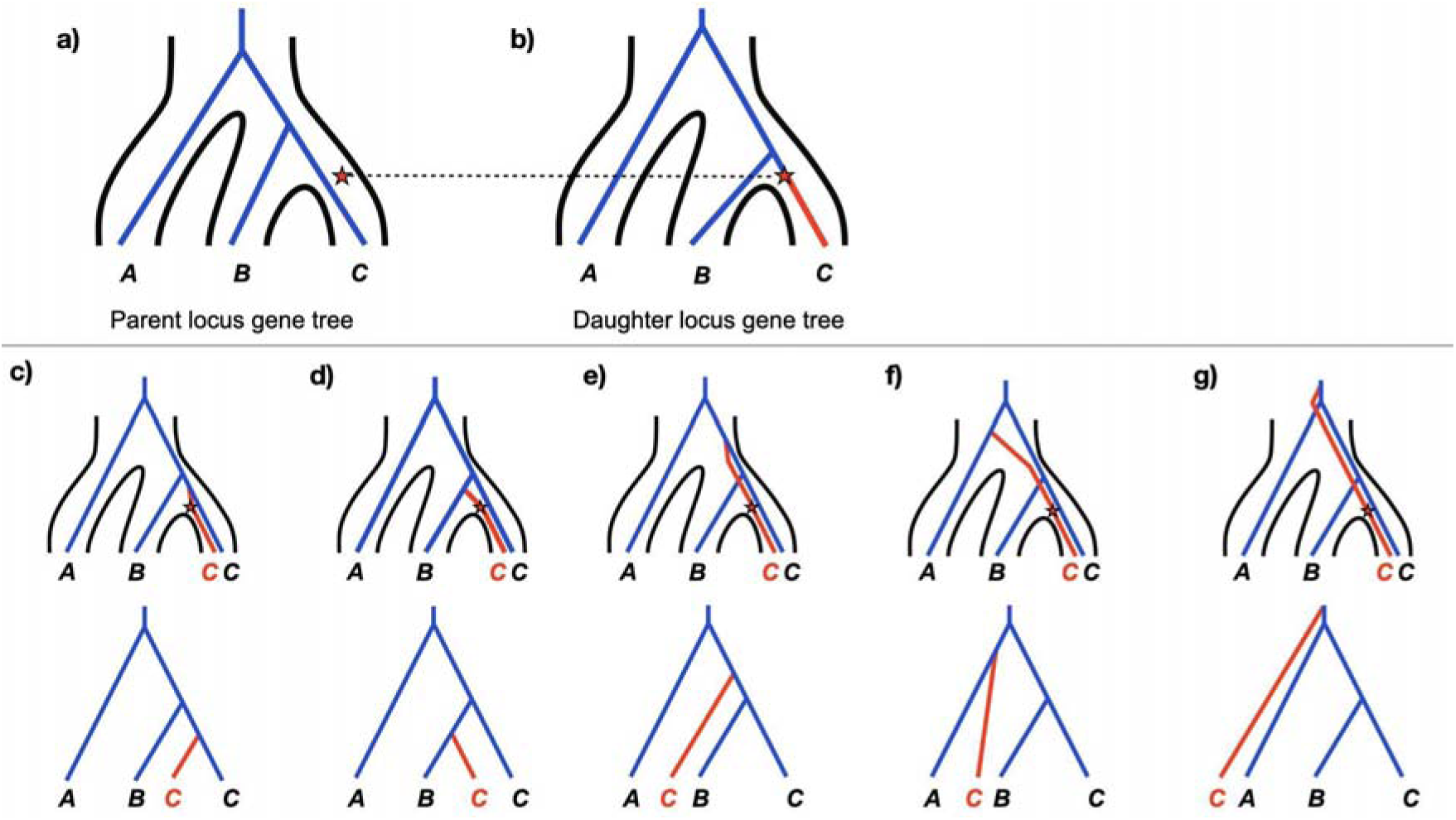
The MSC-DL model generates full gene trees. a) A gene tree at the parent locus (blue) is drawn from the species tree (outlined in black) via the multispecies coalescent (MSC) model. A duplication event (red star) is drawn from a birth-death model, in this case occurring in the population ancestral to species B and C. b) A gene tree at the daughter locus is drawn from the MSC. The duplication is placed on this gene tree on the branch and at the time at which it occurred in the species tree (shown as a dotted line), choosing one of the two possible gene tree branches (B or C) at random. The inserted DNA now takes on the gene tree branch labeled in red. c)-g) Possible ways that the parent and daughter loci coalesce to form the full gene tree. In all panels the star and red branch below it come from the daughter gene tree, and the blue tree comes from the parent gene tree. The coalescent process joining the two trees is shown as a red branch above the star. The trees in the bottom row are simply unfolded views of the full gene trees.

In sum, the model is comprised of three coalescent processes: one that describes gene trees at the parent locus, one that describes gene trees at the daughter locus (or loci), and one that joins parent and daughter gene trees.

### Complex coalescence: ILS, unsorted homologs, and copy-number hemiplasy

The process we are modeling—the interaction of duplication, loss, and coalescence—can be quite complex, leading to both conceptual and linguistic confusion. In this section, we briefly attempt to clarify some important terms that have been used in different ways by different researchers. While we try to avoid introducing new language, we do want to clearly define how terms are being used in this paper and their relation to previous uses.

#### Incomplete lineage sorting (ILS)

ILS occurs when two or more lineages do not coalesce in the population in which they first co-exist. When this happens, these lineages enter an ancestral population, at which time they may again coalesce, either with each other or with additional lineages that are present. If ILS results in lineages from the original population coalescing first with a lineage from a different population, gene trees will be discordant with the species tree. This process can result in discordant gene trees at the parent and daughter loci considered here.

Importantly for the MSC-DL model, the probability of ILS for two lineages is normally expressed as *e^T^*, where *T* represents the length of time during which the lineages can coalesce in the same population, and is usually taken as the length of the relevant branch in the species tree (e.g. Rosenberg 2002; Bryant and Hahn 2020). However, when considering ILS between a pair of duplicates, the amount of time available for coalescence before ILS has occurred is limited to the time between the duplication event and the next speciation event (going backward in time). This time is always shorter than the length of the entire branch of the species tree on which the duplication occurred, which can lead to ILS even when all branches of the species tree are long. The importance of this distinction will hopefully become clearer in the next sub-section.

#### Unsorted homology

Gene duplication is fundamentally a process by which an existing sequence in a population at one locus is inserted into another location. Because of this, the inserted DNA at the new daughter locus carries with it a history of coalescence with all the other sequences at the parent locus, and the full gene tree will reflect this history. In other words, all duplication events also involve coalescence events. Mallo et al. (2014) described a common phenomenon that can occur when duplicates coalesce together, which they called “unsorted homology.” The general idea is that standard DL/DTL methods assume that duplication along a lineage also requires the joining of parent and daughter trees along that same lineage; if this does not occur, there is unsorted homology.

As an example, duplication of a gene along the *C* lineage (Figure 1b) is assumed to only lead to gene trees with the two copies in species *C* as sister to one another (Figure 1c). Any alternative joining of these trees will lead DL/DTL methods to incorrect reconciliations, which will in turn lead to incorrect homology assignments (i.e. orthology and paralogy assignments). However, due to the coalescent process, many other trees can be formed (Figure 1d-g). Because of the mismatch between the lineage containing the duplicate and the lineage to which it coalesces in Figures 1d-g, all of these trees would be considered unsorted homologs. Note that, in the scenario shown here—i.e. conditional on the timing of duplication and no coalescence between lineages *B* and *C* at the daughter locus—the trees in Figures 1d and 1e are equiprobable with the topology in Figure 1c, making unsorted homology the most likely outcome.

It should be emphasized that all of the topologies presented in Figure 1 involve only a single duplication and no losses, and neither the parent tree nor the daughter tree is itself discordant due to ILS. In fact, the example in Figure 1d is not technically due to ILS either, since coalescence occurs in the same population of the species tree in which the duplication appeared (the ancestral population of species *B* and *C*).

This example highlights an additional complication of unsorted homology: it can occur with or without ILS, and even when the species tree only contains two species—and therefore neither parent nor daughter gene trees can themselves be discordant. Supplementary Figure S4 shows examples of unsorted homology in a two-species tree with ILS, while Supplementary Figure S5 shows an example without ILS. One can see that there is very little time for coalescence (i.e. *T*) before ILS occurs in Supplementary Figure S4, as it is the timing of duplication relative to speciation, not the length of the branch in the species tree, that determines the probability of ILS. Some approaches to reduce the effects of ILS on reconciliation have required the presence of “apparent duplications” (Lafond et al. 2014) or “observable duplications” (Scornavacca et al. 2011)—essentially, duplication nodes that have more than one copy of the gene in a descendant species. While such approaches do prevent the inference of duplication when none has occurred, as shown by the examples of unsorted homology here (Figure 1d-f, Supplementary Figure S4, Supplementary Figure S5), they do not ensure the correct identification of duplication nodes, nor do they prevent inferences of excess losses.

#### Hemiplasy

Gene trees at any locus in the genome can be discordant with the species tree. In such cases, the term “hemiplasy” (Avise and Robinson 2008) describes the incorrect inference of homoplasy that can arise when mutations occur on a discordant branch of a gene tree (i.e. an internal branch that does not exist in the species tree). In the context of either duplication or loss, such mutations must occur on a discordant branch of a gene tree in order to lead to “copy-number hemiplasy” (CNH). Loss events can cause CNH when they occur either on a discordant parent tree (Supplementary Figure S6a) or a discordant daughter tree (Supplementary Figure S6b). In contrast, because duplication events only occur on daughter gene trees, they can only cause CNH on discordant daughter trees (Supplementary Figure S6c,d).

However, there is also a broader definition of copy-number hemiplasy in use, which includes any duplication or loss that does not fix in all descendant lineages, regardless of underlying gene tree topologies (Rasmussen and Kellis 2012; Wu et al. 2014). While this definition of CNH encompasses the stricter one given above, it also includes cases such as the one shown in Figure 1 (and Supplementary Figure S5), which the stricter definition does not include. Because the duplicative mutation in Figure 1 occurs in the common ancestral population of species *B* and *C*, but was only inherited by lineage *C*, this scenario would be considered to be hemiplasy by the broader definition. A similar event occurs in Supplementary Figure S5, even though there is no ILS, and no gene tree discordance is possible. Importantly, unlike the strict definition of CNH (and unlike unsorted homology), there is not necessarily a genealogical signal left by events under this broader definition of CNH. For instance, if the duplication event in Figure 1a occurred a few generations later—after the split between species *B* and *C*, along a tip lineage—the scenario would not be considered hemiplasy by this broader definition, but would still produce the same range of gene trees (i.e. Figure 1c-g). (This could now involve ILS, however.)

Regardless of the exact definitions and terms used, many of these important processes are not modeled by common simulators (e.g. SimPhy [Mallo et al. 2016] does not simulate either type of hemiplasy) and are not considered by common reconciliation algorithms (e.g. DLCpar assumes that neither type of hemiplasy occurs). The new simulator introduced in the next section, `dupcoal`, simulates all of these phenomena—as does MLMSC-II—while also reporting the occurrence of each in generating the underlying gene trees.

### Simulation using dupcoal

We created a new simulator—`dupcoal`—for generating gene trees under the MSC-DL model. This simulator is needed because it tracks all evolutionary events that occur, a necessary step in evaluating the performance of `reconcILS.` Our algorithm is described in detail in Appendix section 2 (for an overview, see Supplementary Figure S7) and all code can be found at https://github.com/meganlsmith/dupcoal.

In general, this algorithm is similar to the updated “MLMSC-II” simulator of Li et al. (2024) and is capable of generating copy-number hemiplasy, along with interactions between coalescence, duplication, and loss not modeled by DL and DTL approaches. However, our algorithm differs in a few ways from those of Li et al. (2021; 2024); these differences are discussed in Appendix section 1.

### Reconciliation using reconcILS

#### Overview of main algorithm

Our reconciliation approach uses nearest-neighbor interchange (NNI) rearrangements on trees to model ILS and coalescence events, and to avoid invoking unnecessary duplications and losses (see Appendix section 3 for details). An NNI move on a rooted tree takes the three lineages defined by a single internal branch, for instance (A,(*B,C*)), and rearranges them into the two alternative trees, either (B,(*A,C*)) or (C,(*B,A*)). Any two bifurcating trees with the same set of labels at the tips can always be transformed one into the other by a series of NNI rearrangements (Robinson 1971).

NNI has been used previously in reconciliation algorithms to correct gene trees that may have been inferred incorrectly (e.g. Chen et al. 2000; Chaudhary et al. 2011; Górecki and Eulenstein 2011; Nguyen et al. 2013). Here, we use NNI because the effects of ILS on gene trees can be modeled as a series of NNI rearrangements (Than and Rosenberg 2013), not because there is necessarily a biological link between NNI and ILS. For instance, for three species undergoing ILS, there are two possible discordant topologies, each of which is one NNI move away from the species tree topology (e.g. Figure 2). For four lineages undergoing ILS, there are 14 possible discordant topologies, each of which are between one and three NNI moves away from the species tree topology. Our algorithm takes advantage of this connection between NNI and ILS in order to reconcile gene trees, but can also accommodate gene tree error and introgression using the same NNI events (see Results).

**Figure 2.**
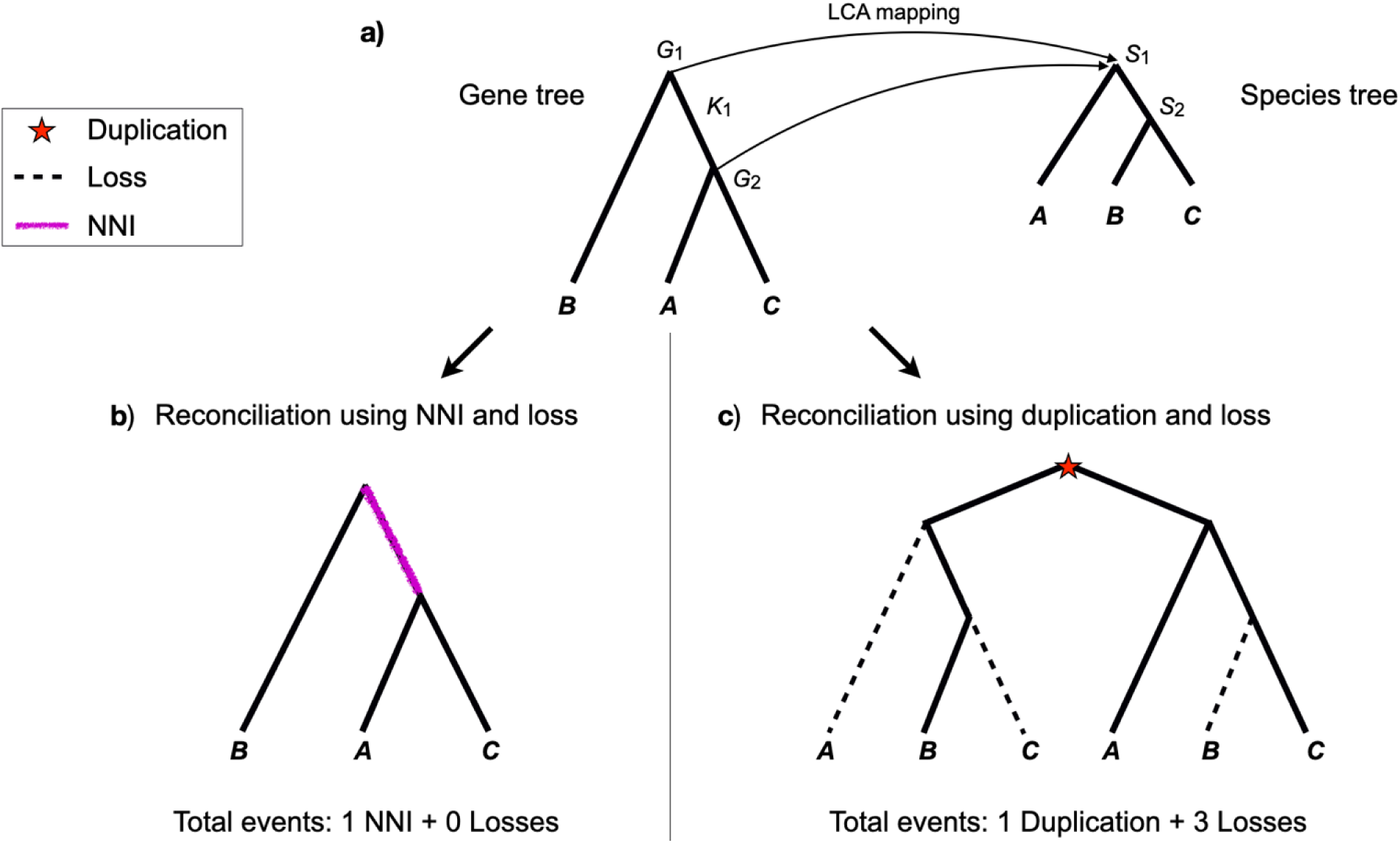
Summary of reconcILS. a) The initial step generates an LCA mapping function between the gene tree and the species tree. Here, nodes and in the gene tree both map to node in the species tree; we therefore refer to as a multiply mapped node. Additionally, and together define a discordant branch, , in the gene tree. For clarity, all mappings involving tip nodes have been excluded. b) The algorithm attempts to reconcile the discordant branch using NNI and loss. Here, reconciliation can be carried out with 1 NNI and 0 losses. c) The algorithm attempts to reconcile the discordant branch using duplication and loss. Here, reconciliation can be carried out with 1 duplication and 3 losses. `reconcILS` must decide between the reconciliations in b) and c) for every discordant branch.

An overview of `reconcILS` is given in Algorithm 1 (see Appendix). Our approach combines multiple sub-approaches that are commonly used in DL reconciliation and other phylogenetic inferences. As a first step, we use the last common ancestor (LCA) algorithm (Zmasek and Eddy 2001) to create a mapping function between each node in a gene tree, , and nodes in a species tree, (Figure 2a). We refer to each of these pairwise relationships (arrows in Figure 2a) as “mappings” from here on.

Unlike in standard DL methods, the LCA mapping here does not represent a final reconciliation. Instead, this step identifies cases in which multiple nodes in the gene tree map to an identical node in the species tree. For instance, in Figure 2a nodes *G*_l_ and *G*_2_ in the gene tree both map to node *S*_l_ in the species tree. We therefore call *S*_l_ a “multiply mapped” node. Additionally, we refer to the branch in the gene tree bracketed by nodes *G*_l_ and *G*_2_ as a “discordant” branch (labeled *K*_l_ in Figure 2a); such branches do not exist in the species tree. The focus of our method is in determining whether discordant branches should be resolved via NNI and loss (Figure 2b) or via duplication and loss (Figure 2c). Choosing among events (NNI, duplication, and loss) is carried out greedily for each branch, with no consideration of a globally optimal set of events among all discordant branches in the gene tree. Consequently, after choosing a set of events for one branch (given specified costs), the process repeats itself until there are no multiply mapped nodes and no discordant branches. Duplications implied by multiply mapped tip nodes in the species tree are inferred via standard DL methods; inferences of losses are explained in Appendix section 3.

As with all parsimony-based reconciliations, `reconcILS` requires a cost for each event type: one for duplications (*C_D_*), one for losses (*C_L_*), and one for NNIs (*C_N_*). Reconciliations can be very sensitive to costs, such that maximum parsimony solutions under one set of costs can be very different than under another set of costs. Although there are few ways to determine costs independently, there are some approaches for finding the Pareto-optimal costs within a reconciliation context (e.g. Libeskind-Hadas et al. 2014; Mawhorter et al. 2019). In the Appendix we discuss additional methods for determining costs, while also demonstrating the robustness of `reconcILS` to the exact choice of costs. Here, as in most commonly used reconciliation algorithms, we allow the user to set costs. The default costs within `reconcILS` are set to *C_D_ = 1.1*, *C_L_ = 1*, and *C_N_ = 1*.

#### Labeling a reconciled gene tree

A full reconciliation of a gene tree, *G*, and species tree, *S*, will have reconciled all multiply mapped nodes, leaving no discordant branches. The total number of duplications, losses, and NNIs required for this reconciliation is the sum of each event type across all branches; these numbers are reported by `reconcILS`. We also aim to label the gene tree with these events—identifying where each event occurred in history—as this is an important output of reconciliation algorithms. As `reconcILS` traverses the species tree finding optimal local reconciliations for each branch, it also records the number and location of all events. Labeling therefore consists of placing these events on the gene tree. Traditional DL reconciliation algorithms label gene tree nodes as either *D* (for duplication) or *S* (for speciation); they may also label the location of losses, *L*. Here, we additionally aim to label the NNI events induced by ILS (N).

Importantly, there is no standard way to label ILS events on a tree, largely because there is no exact map between gene tree nodes and species tree nodes under ILS. A branch undergoes NNI (and ILS), not a node, so it is not clear which node flanking such a branch should be labelled. This labeling problem therefore differs fundamentally from labeling under DL and DTL algorithms because of the lack of a one-to-one map between nodes in the two trees. In the Appendix we explain the labeling used by `reconcILS:` we have created our own labeling system for ILS, one that we think is both understandable and easy to work with. Supplementary Figure S8 shows examples of this labeling for some of the trees in Figure 1.

## Results

### Accuracy and advantages of dupcoal

We assessed the accuracy of `dupcoal` by first simulating 1000 gene trees along a 40-tip species tree with no duplication and no loss (Supplementary Figure S9). These simulations therefore only model ILS as a cause of discordance. As one measure of accuracy, we compared the degree of discordance in these simulations to ones carried out using the same species tree with the simulation program SimPhy (after appropriate branch-length scaling; see Appendix section 2). We found a very high correlation between concordance factors on each branch calculated from gene trees produced by the two methods (Spearman’s *ρ*=0.98, *P*=6.2 x 10^-29^). In a separate study, we have also shown that the gene tree probabilities and branch lengths produced by `dupcoal` match theoretical predictions (Mishra and Hahn 2026). Here, we also carried out simulations in `dupcoal` with both duplication and loss, again comparing our results to SimPhy (after scaling duplication and loss rates appropriately). Again, the average number of duplications and losses per simulated gene tree (*n*=1000) were highly similar and not significantly different between the two methods: 3.71 vs. 3.82 duplications and 2.92 vs. 2.93 losses for `dupcoal` and SimPhy, respectively (*P*=0.43 for duplications and *P*=0.82 for losses; t-tests), suggesting that we are accurately simulating coalescence, duplications, and losses in `dupcoal`.

To highlight additional features contained within `dupcoal`, we also tracked the number of simulated gene trees in which copy-number hemiplasy occurred (this is a standard output reported by `dupcoal`). Out of 1000 gene trees simulated on the 40-tip tree with duplication, loss, and coalescence, 493 contained at least one (strict) hemiplasy event. As a straightforward way to demonstrate the effect of such events in a simulation in which they have occurred, we separately inferred the number of duplications and losses on gene trees with and without copy-number hemiplasy using the DL method implemented in ETE3 (Huerta-Cepas et al. 2016). While ETE3 does not model any type of ILS (see next section), this contrast shows the effects of hemiplasy above and beyond the effects of ILS. As expected, both the number of inferred duplications (2.23 excess duplications; *P*=3.72 x 10^-8^, t-test) and the number of inferred losses (6.89 excess losses; P=4.83 x 10^-9^, t-test) were higher among the trees containing hemiplasy. These results emphasize the importance of having a simulation engine that can simulate hemiplasy.

### Accuracy and advantages of **reconcILS**

As a simple test of DL reconciliation algorithms—which do not allow for ILS—we again used the DL method implemented in ETE3 (Huerta-Cepas et al. 2016). We initially ran simulations in `dupcoal` without duplication or loss (i.e. ILS only); this allows us to see incorrectly inferred duplications and losses more easily. As expected, on a species tree with only 3 tips, DL-reconciliation always inferred 1 extra duplication and 3 extra losses on gene trees with discordance due solely to ILS (Supplementary Figure S10). On a larger species tree with 40 tips, DL-reconciliation inferred large excesses of duplications and losses when only ILS was present (9.6 duplications and 34.8 losses per gene tree, on average). On datasets simulated on a 40-tip tree with duplication, loss, and ILS, the accuracy of DL-reconciliation remains low for duplications (mean absolute error [MAE]=8.5) and losses (MAE=33.6) (Supplementary Figure S11). These results suggest that the coalescent process is a major source of error for DL reconciliation methods.

In contrast, `reconcILS` accurately infers the numbers of duplications and losses (Figure 3a and 3b), as well as ILS events (Figure 3c) on the 40-tip tree. The correlations between inferred and simulated duplications and losses were high (*ρ*=0.95 and 0.63, respectively), as was the accuracy (MAE=0.57 and 1.37, respectively; Supplementary Table S1). We also observed high accuracy in the inferred number of NNI events needed to reconcile a gene tree (*ρ*=0.95; MAE=2.06; Supplementary Table S1). It should be noted that all inferences here are carried out with the true gene trees, as we have not included any (error-prone) steps to infer the trees from sequence data; later, we relax this assumption.

**Figure 3.**
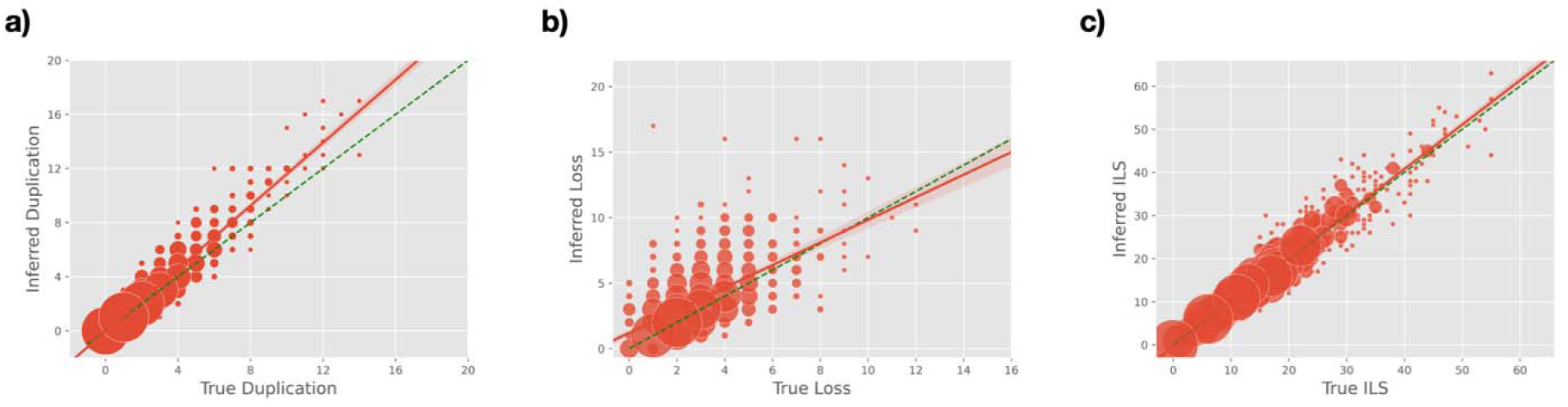
Accuracy of reconcILS on simulated data. 1000 gene trees were simulated on a 40-tip species tree. a) Number of inferred versus true duplication events. b) Number of inferred versus true loss events. c) Number of inferred versus true ILS (i.e. NNI) events. Circle size is proportional to the number of reconciliations with each value, with jitter applied for clarity. Orange solid line represents best-fit regression (as calculated in Seaborn; Waskom 2021), plus confidence intervals. The dashed line represents y=x. See Supplementary Figure S11 for results from ETE3 on the same data.

We also evaluated the performance of `reconcILS` on data simulated from the same 40-tip species tree using SimPhy. Again, the accuracy of `reconcILS` was very high, similar to the accuracy on the `dupcoal` simulations (MAE=0.63 and 1.17 for duplication and loss, respectively). Note that we cannot calculate the accuracy of NNI inference on these trees, as SimPhy does not report the number of NNIs per gene tree. However, the number of “extra lineages” per tree (Maddison 1997) is reported by SimPhy and should be correlated with the number of NNIs; indeed, the correlation between the number of NNIs inferred by `reconcILS` and the number extra lineages was very high (*ρ*=0.94).

As we did with ETE3 in the previous section, we also contrasted the number of duplications and losses inferred by `reconcILS` on simulated gene trees with and without copy-number hemiplasy (using default costs). Unfortunately, `reconcILS` also does not deal perfectly with strict hemiplasy, though in a slightly different way than ETE3. Rather than large overestimates of both the number of duplications and losses, `reconcILS` appears to generally infer an ILS event as a loss event instead. This occurs because (for instance) a duplication on a discordant daughter gene tree branch will duplicate a non-sister pair of lineages (e.g. *A* and *B*), which `reconcILS` interprets as a duplication and a loss (of *C*, in this case) rather than as a duplication and an NNI event (Supplementary Figure S6c). Consistent with this, we observed a small excess of duplications in the simulations with hemiplasy (0.39 extra duplications, on average; *P*=1.49 x 10^-10^, t-test), but significantly more losses (1.04; *P*=7.03 x 10^-17^) and no difference in NNI events (0.05; P=0.78) compared to simulations that did not have hemiplasy.

For completeness, we compared `reconcILS` to ETE3 under several additional conditions. First, we used the original 40-tip tree topology (Supplementary Figure S9) but with very long species tree branch lengths (x1000) to simulate 1,000 gene trees with duplication and loss, but very little ILS. Under these conditions, inferences of duplication and loss using `reconcILS` were even more accurate (*ρ*=0.98 and 0.86, respectively; MAE=0.09 and 0.32, respectively; Supplementary Table S1). Even in these simulations, very rare ILS events occur when there is little time between duplication and speciation (requiring 0.126 NNIs per tree; Supplementary Table S1), with `reconcILS` still accurately detecting these events (MAE=0.32). As expected under these conditions, DL-reconciliation was highly accurate in its inferences of duplications and losses (*ρ*=0.99 and 0.92, respectively; MAE=0.05 and 0.41, respectively), similar to `reconcILS`. We also repeated our analyses using `reconcILS` and ETE3 on 10 additional, randomly drawn species tree of varying heights (Supplementary Table S2). We compared the two methods by taking the ratio of events inferred by ETE3 to the number of events inferred by `reconcILS`. This ratio varied from 1.84-21.1 for duplications (i.e. ETE3 inferred 1.84x-21.1x as many duplications as `reconcILS`) and from 1.1-26.1 for losses, indicating that `reconcILS` was always more accurate (Supplementary Table S2). As expected, these ratios were strongly negatively correlated with tree height (r=-0.90 and -0.94 for duplications and losses, respectively), since shorter trees necessarily have more ILS (because all trees have 40 species).

We also added error to the trees generated under the original 40-tip tree with duplication, loss, and ILS (see Methods). At the lowest level of error (mean of 5 errors per tree), both `reconcILS` and ETE3 show slight reductions in accuracy (increases in `reconcILS` MAE of 0.14, 0.36, and 1.95 for duplication, loss, and NNI, respectively; Supplementary Table S1). As more error is added, both methods continue to become less accurate. At extreme levels of error (mean of 15 errors per tree), while `reconcILS` shows reduced accuracy compared to the baseline case (MAE=1.95, 7.80, and 9.68 for duplication, loss, and NNI, respectively), ETE3 has much lower accuracy (MAE=15.80 and 66.27 for duplication and loss, respectively; Supplementary Table S1).

In the Appendix (section 3) we carry out further analyses, contrasting `reconcILS` with DLCpar and testing both the effects of different costs and unequal duplication and loss rates on our inferences. In general, DLCpar shows equivalent accuracy to `reconcILS`, but cannot be run on gene trees with more than 25 tips; all `reconcILS` results are robust to the choice of event costs and unequal duplication and loss rates.

### Analysis of primate genomes

To further demonstrate the power and accuracy of `reconcILS`, we applied it to a dataset of gene trees inferred from 23 primate genomes (Vanderpool et al. 2020; Smith et al. 2022). These genomes all come from the suborder Haplorhini, which includes all primates except the lemurs and lorises (suborder Strepsirhini). Among these genomes, we analyzed two sets of gene trees that were previously inferred. First, we considered 1,820 single-copy orthologs. This dataset is not expected to contain any duplicates, though it does contain losses. However, because there are high levels of ILS (and some introgression) among the primates sampled here (Vanderpool et al. 2020), the dataset serves as an ideal comparator for reconciliation approaches with and without ILS. Second, 11,555 trees containing both orthologs and paralogs were used to infer duplications, losses, and ILS among the primates. We used both `reconcILS` and ETE3 to analyze the two datasets, assigning inferred events to each branch of the primate tree.

### Single-copy ortholog dataset

Results obtained from analysis of only the single-copy orthologs are shown graphically in Figure 4a (see Supplementary Figure S12a for numerical values on each branch). For this dataset, the average computer memory required per tree was 10.8 MB, while the average analysis time was 1.01 seconds (<31 minutes to analyze all 1,820 gene trees). While this is a somewhat contrived example—a set of only single-copy genes would not be analyzed on its own using reconciliation methods—we think the comparison with the full dataset is highly informative. For instance, the pattern that stands out from this graph is that DL-reconciliation is vastly overestimating the number of duplications and, especially, the number of losses. Because ETE3 does not model ILS, all such events must be reconciled by one duplication (placed above the discordant branch) and three losses (placed below the discordant branch; Hahn 2007). We can see this effect in the data by looking at the correlation between the gene concordance factor (gCF) estimated for a branch and the number of duplications inferred on its parent branch. Gene concordance factors quantify the fraction of loci that contain the same branch as the species tree; when single-copy genes are analyzed, low gCFs can indicate discordance due to ILS or other factors, like gene tree estimation error (Baum 2007; Lanfear and Hahn 2024). The correlation between gCFs for these data (estimated in Smith et al. 2022) and duplications inferred by ETE3 on parent branches is ρ=-0.56. Lower gCFs lead to higher numbers of duplicates, as expected when ILS is not accounted for—this problem is not unique to ETE3, but should be expected for all standard DL and DTL approaches. Such errors can lead to incorrect inferences of bursts of duplication and loss.

**Figure 4.**
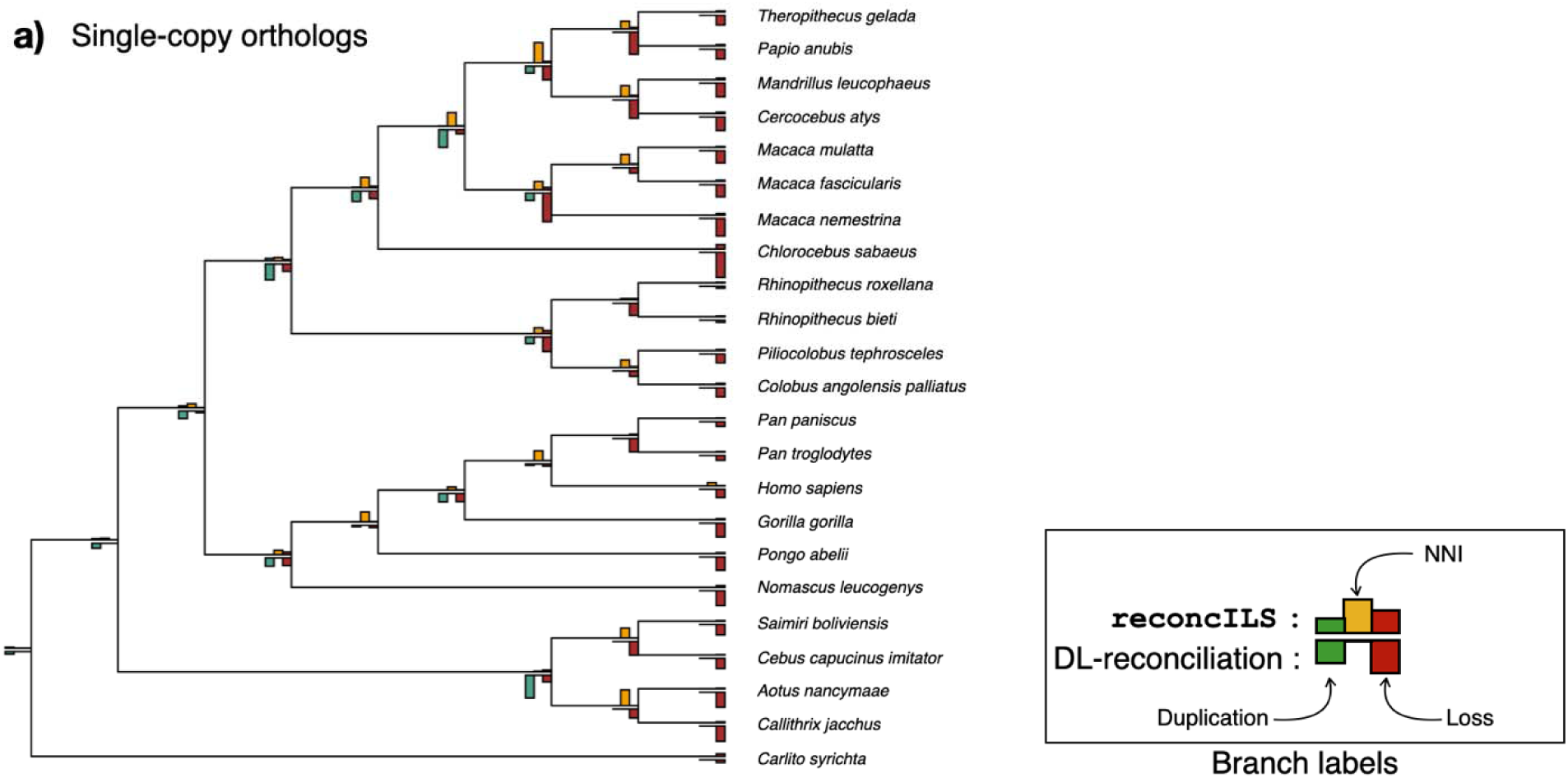
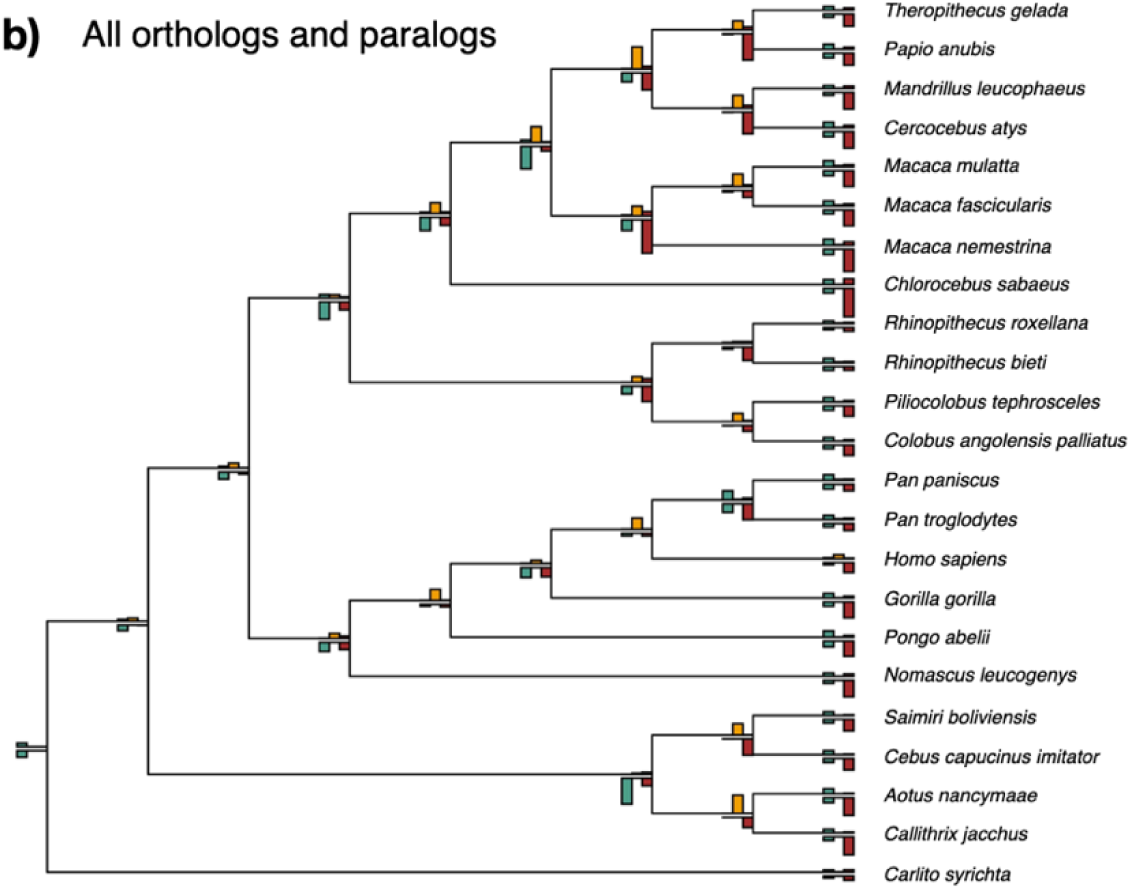
Results from primates. a) Analysis of 1,820 single-copy orthologs. Both `reconcILS` and ETE3 (a DL-reconciliation method) were used to analyze the data, with duplication, NNI, and loss events mapped to each branch of the species tree. On each branch, the bar plots show results from `reconcILS` on top, with results from ETE3 mirrored below. ETE3 cannot infer NNI events, so this bar is always set to 0. b) Analysis of 11,555 gene trees containing both orthologs and paralogs. Results are presented in the same manner as in pane a. However, note that the height of all bars is normalized within a panel, so that heights of bars within a tree can be compared among branches, but the heights of bars between the trees cannot be compared. Supplementary Figure S12 shows the absolute counts of all events from both panels and Supplementary Figure S13 shows species tree with branch lengths in substitution units rather than as a cladogram, as shown here. Trees were drawn with ggtree (Yu et al. 2017).

Using the same reasoning, we can assess the accuracy of `reconcILS` by examining the correlation between NNI events inferred by our algorithm and gCF values for single-copy genes. High NNI counts indicate more ILS, and therefore this measure should be negatively correlated with gCF values for the same branch. Indeed, the correlation between NNI counts and gCF values is *ρ*=-0.83, demonstrating the high accuracy of `reconcILS` in quantifying ILS (or in correcting for gene tree error). However, `reconcILS` does sometimes infer a small number of duplications (Figure 4a; Supplementary Figure S12a). While it is possible that some of the single-copy genes we analyzed represent so-called pseudoorthologs—and therefore do contain actual duplication events—these are expected to be rare (Smith and Hahn 2022). Instead, these false duplicates are likely due to ILS among more than four lineages. Recall that, although our algorithm has been developed to account for rearrangements due to the coalescent process, ILS among many lineages can result in gene trees that are many NNI moves away from the species tree. For instance, ILS among four lineages can result in gene trees between one and three NNIs away from the species tree, though in these cases the NNI solution will still be favored over duplication and loss. However, ILS among more than four lineages will produce trees that are even further away. Therefore, multiple NNI events may sometimes simply be equally or more costly than duplication and loss, even when ILS is the true cause of discordance (`reconcILS` reports the duplication+loss solution to a reconciliation when there are equally optimal reconciliations).

### Full gene tree dataset

We also examined patterns of ILS, duplication, and loss for the entire primate dataset. Given that these data consist of more and larger gene trees than the single-copy dataset, the average computer memory required per tree was slightly higher (17.1 MB), while the average analysis time was also higher per tree (5.05 seconds; ∼16 hours to analyze all 11,555 gene trees). The results from the full set of gene trees (Figure 4b; Supplementary Figure S12b) largely mirror the results from the single-copy orthologs (which are a subset of the entire dataset). Part of the apparent similarity is simply due to normalizing the two datasets separately, but several non-graphical factors also contribute to the pattern. First, although there are many true duplicates in the full dataset, the vast majority of gene trees only have events on tip branches: in 7,693 out of 11,555 gene trees all duplicates are specific to one species (Smith et al. 2022). Another *∼*1,000 trees have duplicates specific to two species. Thus, there are relatively few trees with duplicates on internal branches even in the full dataset.

Second, errors in inferring duplications and losses due to the coalescent process still dominate the results from standard DL reconciliation (i.e. ETE3), possibly even more so than in the single-copy dataset. Because ILS (and introgression) can affect all gene trees, trees with duplicates can experience twice as much discordance since there are twice as many branches. One example of this can be seen among the Papionini (macaques, baboons, geladas, and mandrills): ETE3 has to infer *>*6,000 losses on each of the tip-branches in this clade, and more than 10,000 losses on most of the internal branches (Figure 4b; Supplementary Figure S12b). There are also many thousands of duplicates inferred on internal branches of this subclade, though these species have the same number of total protein-coding genes as all the others analyzed here (Vanderpool et al. 2020). Although the inferred duplications and losses in this group were also relatively higher in the single-copy dataset, it is clear that even a small number of true duplicates can have an outsized effect on methods that cannot account for ILS.

The analysis using `reconcILS` is also highly similar between the two datasets, again largely because most duplicates are confined to tip branches. `reconcILS` is able to deal with discordance due to coalescence, even in the presence of duplication: the correlation between NNI events inferred on each branch of the species tree from the single-copy orthologs and the full dataset is *ρ=*0.99. Although the numbers of events are of course larger with the larger dataset (Supplementary Figure S12b), the effect of ILS on any given branch of the species tree is experienced by all gene trees. Given its ability to deal with ILS, duplication events across the primate tree are more reliably inferred with `reconcILS`.

## Discussion

We have introduced a new reconciliation algorithm and associated software package (`reconcILS`) that considers discordance due to duplication, loss, and coalescence. To better demonstrate the accuracy of our method, we have also developed a new simulation program (`dupcoal`) that generates gene trees while allowing these processes to interact under the MSC-DL model. Our results demonstrate that `reconcILS` is a fast and accurate approach to gene tree reconciliation, one that offers multiple advantages over alternative approaches, even those that also consider ILS (e.g. DLCpar; Wu et al. 2014). Nevertheless, there are still several caveats to consider, as well as future directions to explore.

Despite our demonstration of good performance in practice, our method is heuristic: we have no theoretical proof that our method finds most parsimonious reconciliations. While such proofs often require strict assumptions—e.g. the proofs for DL algorithms assume there is no ILS—they are very helpful in providing confidence under the specified conditions. However, given the high accuracy of `reconcILS` demonstrated with both simulations and empirical data from single-copy genes, we hope the heuristic nature of our algorithm is not a barrier to its utility. Future theoretical work on this or similar approaches using NNIs (e.g. Górecki and Eulenstein 2011) would of course be of benefit to the field. Under any such model, it must also be recognized that errors in gene tree reconstruction will have a detrimental effect on the results (e.g. Supplementary Table 1). Both theoretical work and new algorithms should take such limits into account.

As mentioned earlier, there is often ambiguity in how to label nodes in gene trees affected by ILS and duplication (e.g. Supplementary Figure S8). This is due in part to the nature of ILS—it is not clear which node to label because there is not a one-to-one map between gene tree nodes and species tree nodes—and in part to the intrinsic lack of information contained only within tree topologies when coalescence and duplication interact. The difficulty in labeling is not confined to this work: DLCpar does not label nodes, Notung labels some ambiguous nodes that could be due to either duplication or ILS, and RecPhyloXML, a format for representing reconciled trees, does not include a way to label ILS events at all (Duchemin et al. 2018).

Nevertheless, labeling gene trees serves multiple important functions, and is often interpreted as biological truth. For example, multiple methods use inferred gene tree labels to identify orthologous genes (genes related by speciation events; Fitch 1970). The general approach taken is to label nodes using reconciliation or similar approaches, treating any node not labeled with a *D* as a speciation node; genes whose common ancestor is a speciation node are subsequently defined as orthologs. Both standard DL reconciliation and so-called “overlap” methods (e.g. Huerta-Cepas et al. 2007; Emms and Kelly 2019) will consistently label the top-most node flanking a discordant branch with a duplication event. This means that, in cases like the one shown in Supplementary Figure 8b, these methods will label the wrong node with a *D*, and subsequently identify the wrong pair of genes as orthologs. (Supplementary Figure 8c would be labeled correctly; recall that this outcome has an equal probability to the one in panel b in the scenario considered here.) Cases such as these have been referred to as “unsorted homologs” (Mallo et al. 2014), and will occur any time duplication and coalescence interact, either before or after a speciation event. They can also occur even if only two species are analyzed (e.g. Supplementary Figures S4 and S5), which means that approaches that simply collapse short branches to avoid ILS will also be affected. Currently `reconcILS` only labels one node with an *N*, though this could also be interpreted as labeling the branch above it. In the future we could allow for multiple possible labelings in order to emphasize the inherent ambiguity of some labels.

The algorithm used by `reconcILS` deals, in part, with an additional interaction between duplication, loss, and coalescence: hemiplasy. Hemiplasy is the phenomenon by which a single mutational event on a discordant gene tree will be interpreted as multiple events (“homoplasy”) when reconciled to the species tree (Avise and Robinson 2008; Hahn and Nakhleh 2016). In analyses of nucleotide substitutions, generic binary traits, and even continuous traits, hemiplasy requires that mutations have occurred on a discordant branch of a gene tree (Mendes and Hahn 2016; Guerrero and Hahn 2018; Hibbins et al. 2023). In the context of duplication, such mutations would have to occur on a discordant branch of a daughter gene tree to lead to copy-number hemiplasy (Supplementary Figure S6c); for loss, mutations can occur on discordant branches of either the parent gene tree or the daughter gene tree (Supplementary Figure S6a,b). The simulation engine introduced here, `dupcoal`, simulates hemiplasy events and reports any gene trees affected. Our results show that these events lead to increased error in reconciliation, and are therefore an important process to include. Interestingly, such events have little effect on species tree inference when paralogs are used for this purpose (Li et al. 2024).

Unfortunately, strict copy-number hemiplasy events as defined above are still not accurately inferred by `reconcILS`: we have found that our algorithm consistently turns ILS into losses when hemiplasy occurs. While the total number of events does not seem to increase, it is not ideal that ILS is mistaken for losses. However, `reconcILS` does deal accurately with copy-number hemiplasy under the broader definition in use (see section on “Complex coalescence…”), and `dupcoal` also simulates and records such events. In contrast, DLCpar assumes that this type of hemiplasy does not occur, and SimPhy does not simulate copy-number hemiplasy at all.

The algorithm used by `reconcILS` deals with ILS differently than does DLCpar. While DLCpar treats ILS as an event, `reconcILS` considers NNIs to be tree operations induced by ILS. It is not clear that either approach should necessarily be preferred on first principles. In simple cases involving three species and a single discordant branch (e.g. Figure 2), counting ILS or NNIs are more or less equivalent, though it should be noted that the process of ILS can also lead to concordant trees and is not synonymous with discordance. However, in cases with more than one discordant branch there is no simple relationship between the two. For instance, in a 4-tip tree undergoing ILS, lineages may undergo ILS across two branches, but result in between 0 and 3 NNI events (Than and Rosenberg 2013). We think that directly considering NNIs—as is done in `reconcILS`—is a natural way to express the events needed to reconcile a gene tree with a species tree, including discordance due to gene tree error. In fact, NNI and other moves have been used previously in reconciliation algorithms to correct gene trees that may have been inferred incorrectly (e.g. Chen et al. 2000; Chaudhary et al. 2011; Górecki and Eulenstein 2011; Nguyen et al. 2013). Though these algorithms minimize DL reconciliation costs using NNIs, they could also have been used within `reconcILS` for each local reconciliation step. In addition, the effects of introgression can be modeled as a series of NNIs (Mishra et al. 2026). This implies that, in cases with introgression (like primates; Vanderpool et al. 2020), using `reconcILS` again helps to avoid inferring excess duplication and losses, though clearly NNIs cannot be directly interpreted as indicating ILS. Regardless of the exact algorithm used, the utility of such methods also provides a motivation for using `reconcILS` to obtain more accurate inferences of duplications and losses in the presence of multiple processes, not just ILS.

There are also several extensions that could be made to `dupcoal`, our new simulation engine. Currently, `dupcoal` draws a daughter gene tree from the multispecies coalescent (MSC) either matching the parent tree (fully linked) or independently from the parent tree (completely unlinked). It is straightforward, however, to draw the daughter tree at a given recombination distance from the parent tree, allowing for partial linkage. In addition, though the software currently draws gene trees from the MSC, we could also include introgression as a source of discordance by drawing trees from the multispecies network coalescent (Meng and Kubatko 2009; Yu et al. 2011). In this case, users would simply input a species network to simulate from, rather than a species tree. We should also note the differences between the current version of `dupcoal` and alternative simulators. As mentioned previously, SimPhy (Mallo et al. 2016) does not simulate hemiplasy of any kind, as it uses the same three-tree model of Rasmussen and Kellis (2012) and Wu et al. (2014). However, this method does include gene transfers, which are not simulated by `dupcoal`. The MLMSC-II simulator of Li et al. (2024) simulates hemiplasy under a similar model to the one used here; `dupcoal` is therefore most similar to this method, with a few differences (see Appendix section 1). Despite these minor differences, both methods are capable of generating a wide variety of gene trees impacted by duplication, loss, coalescence, and the interactions of these processes. Unlike MLMSC-II, `dupcoal` records the exact counts of several different event types, a necessity when evaluating the performance of reconciliation approaches.

Finally, we envision multiple ways in which the `reconcILS` algorithm could be used in conjunction with other methods. For instance, ASTRAL-Pro (Zhang et al. 2020) infers a species tree from gene trees containing paralogs by first labeling nodes in each gene tree as either duplication or speciation. The authors of this software rightfully recognize the problems with labeling discussed here, but nevertheless the labeling step might benefit from methods that include ILS. Likewise, GRAMPA (Thomas et al. 2017) allows for reconciliation of polyploidy events, but does not include coalescence as a source of discordance; its algorithm could easily accommodate this extension. There are also multiple approaches that can be used to correct gene trees via reconciliation (e.g. El-Mabrouk and Ouangraoua 2017; Christensen et al. 2020). These and many other tasks might benefit from approaches that model duplication, loss, and coalescence in a single framework.

## Methods

### dupcoal

The simulator `dupcoal` is written in Python 3 and uses functions from DendroPy (Sukumaran and Holder 2010) to generate trees under the MSC and to manipulate parent and daughter trees. It also uses functions from NumPy (Harris et al. 2020) to draw coalescent times, duplication times, and loss times from exponential distributions. Supplementary Figure S7 gives and overview of the simulation process; further details on implementation and testing can be found in Appendix section 2.

### reconcILS

The reconciliation software `reconcILS` is written in Python 3 and uses multiple packages for tree handling and data plotting; software and required packages can be installed with pip or Conda. The only required inputs to the software are a species tree and at least one gene tree. Details on the algorithm, implementation, and further testing of `reconcILS` can be found in Appendix section 3.

### Primate data

We used two sets of gene trees that were previously inferred in Smith et al. (2022) using IQ-TREE2 (Minh et al. 2020), based on data from Vanderpool et al. (2020). The first set consists of 1,820 single-copy orthologs; this set was denoted as “Single-copy clusters” in Smith et al. (2022). The second set consists of 11,555 trees containing both orthologs and paralogs; this set was denoted “All paralogs” in Smith et al. (2022). We randomly resolved all polytomies. When using `reconcILS`, costs were set to *C_D_ = 1.1*, *C_L_ = 1*, and *C_N_ = 1*.

## Acknowledgements

We thank Yao-ban Chan for many helpful conversations about the MLMSC and MSC-DL models, Jessica Wu for answering questions about DLCpar, and Ben Fulton for tips on best programming practices. Reviewers of the manuscript also provided many constructive comments. This work was supported by NSF grant DEB-1936187 and DBI-2146866 to MWH.

## Data availability

`reconcILS` is available at https://github.com/smishra677/reconcILS. `dupcoal` is available at https://github.com/meganlsmith/dupcoal.

## Appendix

### 1. Similarities and differences between the MSC-DL and MLMSC models, algorithms, and outputs

Here we address the conceptual and algorithmic differences between two models: the MSC-DL (introduced here; see also Mishra and Hahn 2026) and the MLMSC (Li et al. 2021, 2024) that both consider duplication, loss, and coalescence. While the DLCoal model of Rasmussen and Kellis (2012) also considers all three processes, their “three-tree” approach limits the types of coalescent events that take place, and we therefore do not consider it further here. Nonetheless, all three of these models are more complete descriptions of the processes that generate gene trees than are models that consider only duplication and loss.

Both the MSC-DL and the MLMSC models treat each individual locus as evolving via the multispecies coalescent model, in which discordance can arise only via incomplete lineage sorting. Both models also further describe how relationships among these loci can be generated via duplication and coalescence, and how lineages are removed via losses. However, there are several differences in exactly how these processes are conceptualized and instantiated, differences that we outline here:

- The MSC-DL considers duplications generated directly on the parent trees (step 2 in the `dupcoal` algorithm described below), rather than using the “coalescent-rate” process introduced in Li et al. (2021). This difference in the algorithms should not result in many differences in output, since the gene trees that duplications are placed on are drawn from the same distribution. However, as the parent tree is the source of duplicates in the MSC-DL model, we think that this formulation is more intuitive (and saves a small amount of computation, since we do not have to discard a tree and draw another).
- In MLMSC-II (Li et al. 2024), a rejection step was introduced after the generation of a daughter tree, before the mutation is placed (i.e. the “haplotype forest” rejection step in their algorithm). We do not include such a rejection step, as we always place a duplication on a branch of the first daughter tree drawn (step 3 in the `dupcoal` algorithm). A related minor bookkeeping difference is that a full gene tree is drawn at the daughter locus in the MSC-DL, and not a gene tree truncated at the time of duplication, as in the MLMSC. This means that there are no haplotype forests in the MSC-DL, though the same process of choosing among existing lineages to place the duplication on occurs in both models.
- One possibly important difference between the algorithms occurs when coalescing parent and daughter trees together. The MLMSC algorithm requires that the daughter tree coalesce with its parent before time of origination of the parent locus (see Figure 14 in Li et al. 2021). If it does not coalesce, the duplicate is discarded. The MSC-DL (and `dupcoal`) have no such truncation times at any locus, and therefore all duplicates coalesce and are kept.
- One related conceptual difference between the models is that the MSC-DL model specifies that duplications at daughter loci can occur only in a lineage “that contains genetic material descended from the original parent locus” (point vi in the main text). In other words, duplication events at daughter loci can only duplicate material that itself was duplicated—otherwise, they would not form part of the same full gene tree. As a consequence of this condition and the previous point above, when `dupcoal` coalesces all marginal trees together (step 5), if a duplication fails to coalesce with its parent tree before the time of origination of the parent (which can happen when it itself is a duplicate), it will coalesce it further in the past with the parent of its parent, rather than discarding the duplication as in MLMSC. These so-called “higher-order” duplicates will be quite rare (Li et al. 2024), but may result in some differences between the simulators.

To better understand how the trees generated under the MSC-DL differ from those simulated using the MLMSC, we compared simulations from `dupcoal` and MLMSC-II (Li et al. 2024). We used the species tree from 16 fungal genomes (Butler et al. 2009), which has been used in several papers investigating the impacts of duplication, loss, and ILS (e.g., Rasmussen and Kellis 2012; Li et al. 2024). We considered a duplication rate λ ∈ [0,6] × 10^-10^, a generation time of 0.1 year/ gen, and a loss rate *μ* ∈ [0.5×*λ,λ*]. We set 2*N* = 1 x 10^7^ in MLMSC-II. We also assessed results in which branch lengths were scaled by dividing them by 9 to assess the impacts of shorter branches. We generated 10,000 gene families per condition per program, and we compared the number of duplications and gene copies between the two programs. We retrieved information on the number of observable duplications from the tables output by MLMSC-II and `dupcoal`.

One note on the outputs of the two programs: for each simulated gene family, `dupcoal` outputs the full simulated gene tree, including annotation of parent and daughter copies. Along with the tree, a table for each family reports: the number of duplications, the number of observable duplications (i.e. duplications that are not completely obscured by loss), the number of losses, the number of cases of different types of hemiplasy (see the main text section on “Complex coalescence…”), and the number of ILS events. In contrast, MLMSC-II outputs a full, unannotated gene tree, and the only information output is the number of observed duplications, the number of genes, and the number of species. The runtimes for the two methods were similar, with `dupcoal` being slightly more computationally intensive than MLMSC-II, taking 12.017 cpu seconds to generate 1,000 gene families when λ=μ=2 x 10^-10^, compared to 10.009 cpu seconds for MLMSC-II.

The average number of duplications was similar across programs, with differences increasing with the duplication rate, *λ* (Figure A1A, A1B). MLMSC-II tended to have higher numbers of duplications than `dupcoal`, with only the latter showing a linear relationship between the number of duplications and the duplication rate. The average number of gene copies was very similar across the two programs when *μ* was smaller than *λ* (0.5X; Figure A1C). However, differences were more pronounced when *λ* and *μ* were equal. In MLMSC-II, the average number of gene copies increases with the duplication rate, even when duplication and loss rates are equal (Figure A1D). On the other hand, the average number of gene copies in `dupcoal` remains constant (and equal to the number of species) when duplication and loss rates are equal, regardless of the duplication rate (Figure A1D). Under trees with shorter branches (i.e. smaller effective population sizes), both methods produced constant numbers of gene copies (Figure A1D). We emphasize that the behavior of dupcoal is the expected result. Investigating further, the behavior of MLMSC-II appears to be due to this program automatically re-running all simulations with fewer than four gene copies remaining in the final tree. If we filter dupcoal results to only those with >= 4 tips, we can reproduce the pattern seen in MLMSC-II. This filtering also eliminates the othe minor differences observed across the two programs.

**Figure A1.**
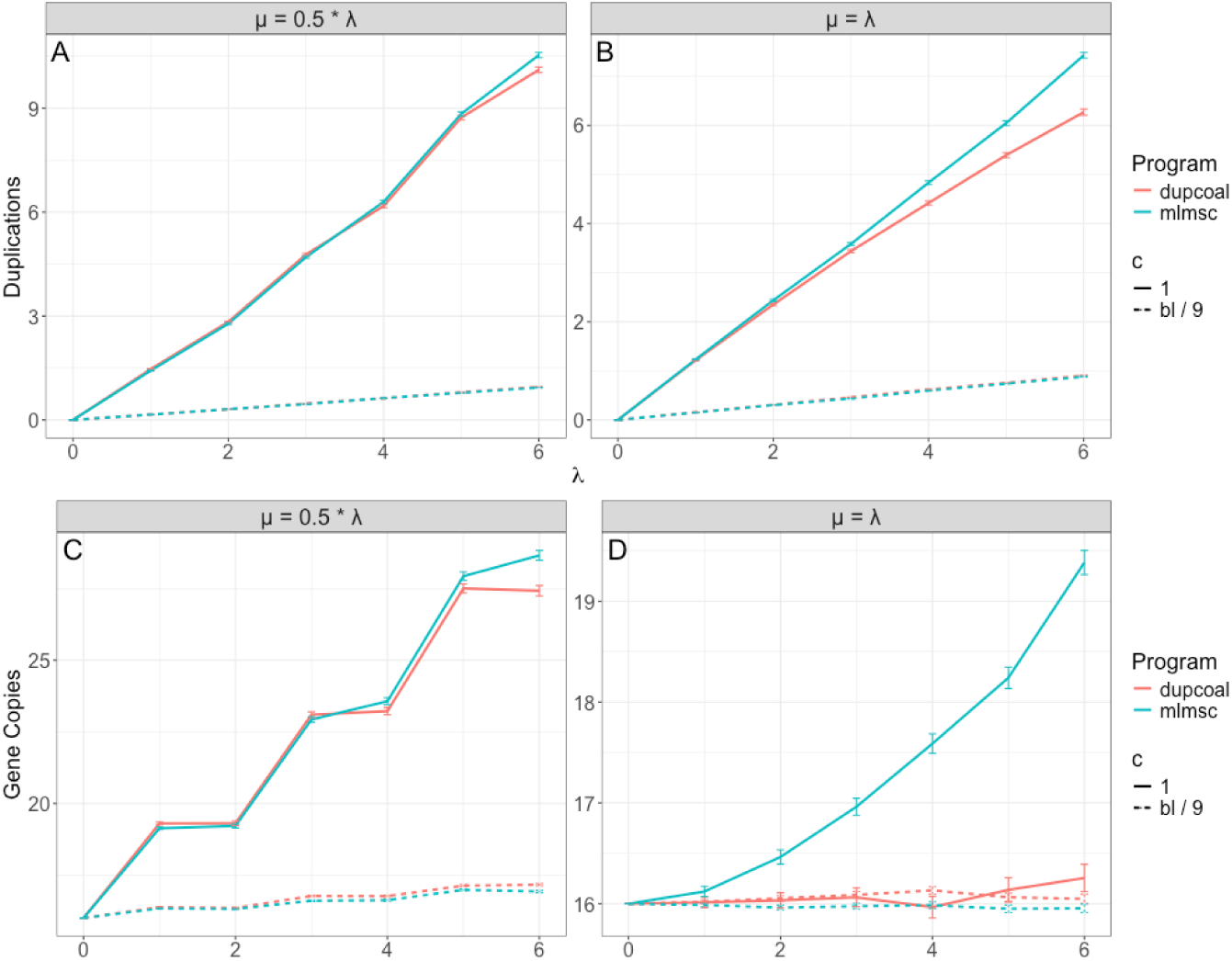
Comparison of results simulated by dupcoal and MLMSC-II. The number of observed duplications (panels A, B) and gene copies (panels C, D) generated under different programs, duplication rates, and loss rates. A) The number of observed duplications across different values of λ when μ = 0.5 * λ. B) The number of observed duplications across different values of λ when μ = λ. C) The number of gene copies across different values of λ when μ = 0.5 * λ. B) The number of gene copies across different values of λ when μ = λ. Average values are plotted, with bars representing standard errors.

### 2. dupcoal—implementation and testing

We created a new simulator—`dupcoal`—for generating gene trees under the MSC-DL model. This simulator is needed because it tracks all evolutionary events that occur, a necessary step in evaluating the performance of `reconcILS`. Our algorithm works as follows (see also Supplementary Figure S7; code can be found at https://github.com/meganlsmith/dupcoal):

1. Generate the gene tree at the parent locus from the species tree according to the MSC model.
2. Simulate duplications along the branches of the parent tree and any existing gene trees (only once for any tree, if returning to this step) at a rate *λ*.
3. For each duplication, generate a daughter tree at a new locus according to the MSC model. Then, place the duplication on the daughter tree by randomly selecting a lineage available at the time of duplication on the appropriate branch of the species tree (Figure 1b). Discard all other lineages from the daughter locus, and truncate the sub-tree at the time of duplication.
4. Return to step 2 and recursively generate new duplications (and subsequent daughter trees) on each new daughter tree. Proceed to step 5 when no additional duplications have been generated on the last daughter tree.
5. Coalesce all trees together. Begin at the present and work backwards in time. First, attempt to coalesce a daughter tree with its parent, which could itself be a duplicated tree. If coalescence fails to occur before the tree originates, attempt to coalesce the daughter tree with the parent of its parent, and so on. Repeat until all trees have been joined together.
6. Generate losses on the full gene family tree at a rate *μ*, truncating gene tree branches below loss events. The result is the final gene family tree.

In general, this algorithm is similar to the updated “MLMSC-II” simulator of Li et al. (2024) and is capable of generating copy-number hemiplasy, along with several other interactions between ILS, duplication, and loss not modeled by DL and DTL approaches. However, our algorithm differs in a few ways from those of Li et al. (2021; 2024); these differences were discussed in the previous section.

While our simulator does not currently include the ability to simulate partially linked duplications, introgression, or horizontal gene transfers, it is straightforward to add these types of events in future iterations of our software. `dupcoal` can currently simulate fully linked and fully unlinked duplicates, and partial linkage simply requires that the topology of the daughter gene tree be correlated with the topology of the parent gene tree (proportionally to recombination distance). Both introgression and transfer can be accommodated by drawing gene trees from a different locus history (e.g. a history of introgression) in step 1 or step 3.

#### Simulated datasets

We simulated gene trees in `dupcoal` along species trees with 40 tips. The species tree topologies were drawn at random using the program ape (Paradis and Schliep 2018), with the “Grafen” method for drawing branch lengths (scaled up by 10); this tree was used as input to both `dupcoal` and SimPhy. The amount of ILS in the base 40-tip tree (Supplementary Figure S9), as measured by gene concordance factors (Minh et al. 2020a), ranged from 28.4 to 99.8; in comparison, the values estimated from the primate tree in Vanderpool et al. (2020) ranged from 27 to 96, so the two are quite similar. Because `dupcoal` specifies branch lengths in coalescent units, while SimPhy uses time, branch lengths were converted by assuming a population size of *N*=10 and a generation time of 0.5 year. Simulations were run on this tree with only ILS (setting duplication and loss rates to λ= 0 and *µ*= 0, respectively). When using non-zero duplication and loss rates, they were scaled to the appropriate branch lengths to make them comparable in `dupcoal` and SimPhy. Comparisons between `dupcoal` and SimPhy with no duplication and loss were carried out by calculating gene concordance factors (gCFs) on the species tree using IQ-TREE2 (Minh et al. 2020b).

### 3. reconcILS—implementation and testing

We created a new reconciliation program—`reconcILS`—that infers the number and timing of duplication, loss, and NNI events from gene trees. It additionally labels nodes on these gene trees as duplications, loss, NNIs, or speciation events. Below is a general description of the algorithm (code can be found at https://github.com/smishra677/reconcILS).

#### Inputs and outputs

Given a rooted, bifurcating species tree, *S*, and a rooted, bifurcating gene tree, *G*, as input, `reconcILS` outputs the number of duplications, losses, and NNIs required to reconcile *G* and *S* given event costs. Inferred events are all also assigned to nodes in the gene tree and tallied on branches of the species tree.

#### Traversing trees and inferring losses

After generating an initial LCA mapping (Zmasek and Eddy 2001) between a gene tree and the species tree, `reconcILS` performs a preorder traversal of the species tree. For each species tree node, the algorithm checks how many gene tree nodes are mapped to it. If there are fewer than 2 mappings, the algorithm asks whether speciation or loss can be inferred. As is standard in reconciliation, species tree nodes mapped by only one gene tree node imply a speciation event at that gene tree node—these do not incur a cost. Nodes mapped by zero gene tree nodes imply at least one loss in the gene tree. Our program asks whether the parent node of a node with 0 mappings itself has 0 mappings. If not, a loss is inferred; if so, no event is recorded, because the loss must have occurred further up the tree. If there are 2 or more mappings the species tree node is multiply mapped, and a function to carry out “local reconciliation” is called. Local reconciliation is described in more detail in the next section, but here can simply be described as determining the events that have generated the multiple mappings to a single species tree node. It should also be noted that if a duplication node is inferred for a gene tree, both sub-trees induced by the duplication are further locally reconciled (Algorithm 1); this means that `reconcILS` can detect cases in which multiple independent losses have occurred along the same lineage (e.g. Figure A4b).

If there are only two adjacent nodes in the gene tree mapped to a node in the species tree (i.e. one discordant branch, as in Figure 2 in the main text), then the method proceeds as described below. However, when there are more than two nodes mapped to an internal species tree node—and therefore more than one discordant branch in the gene tree—we must decide which branch to reconcile with the multiply mapped node first. We have found cases where the order in which the branches are resolved matters, with only one of the choices leading to an optimal reconciliation. Algorithm 2 describes an algorithm for choosing which discordant branch to locally reconcile first. This algorithm calculates the “bipartition cost” of each possible branch to reconcile, in effect efficiently looking at the future costs associated with alternative NNI moves (Figure A2). We calculate the bipartition cost of a branch by examining the two trees induced by an NNI move applied to the branch. For each of these new topologies, we calculate costs using two measures, both of which are applied independently to the two sub-trees extending from the root of these topologies (denoted by the red vertical line in Figure A2). First, we count the number of bipartitions present in these sub-trees but not present in the species tree (“bipartition score” in Figure A2); second, we count the number of lineages missing relative to the node in the species tree the root of the topology is mapped to (“lost cost” in Figure A2). The bipartition cost is the sum of the bipartition scores and lost costs of a single rearranged tree; this step therefore requires eight values to be calculated for each branch considered, four for each of the trees created by NNI. Such calculations are simple and can be carried out quickly, and by choosing the branch with the lowest bipartition cost we can more accurately proceed with reconciliation.

Once reconciliation for a single multiply mapped node in the species tree is carried out (i.e. it is no longer multiply mapped), our algorithm begins again: a new round. In each round we generate an LCA mapping function and continue traversing the species tree, counting gene tree nodes mapped to each species tree node.

**Figure A2.**
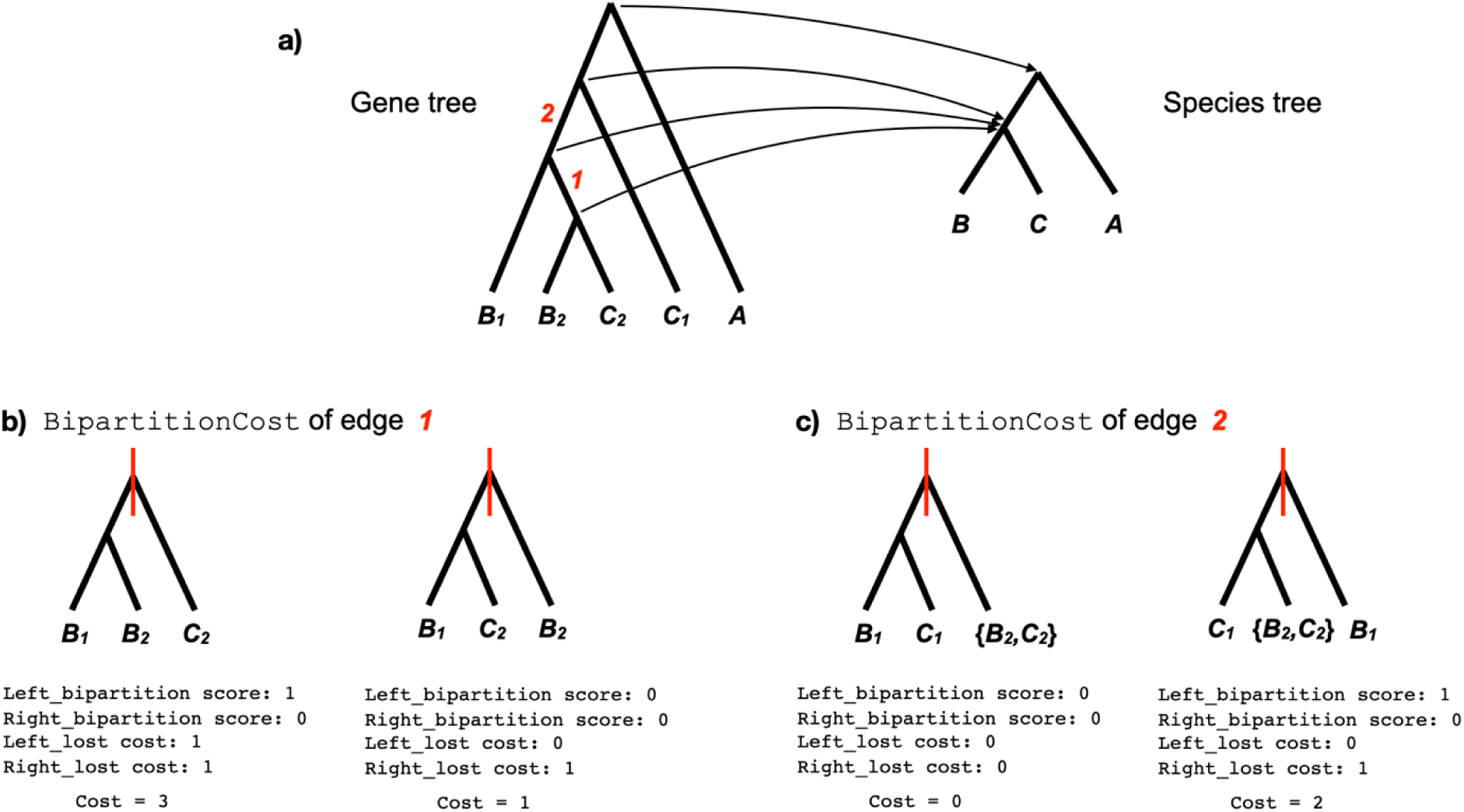
Calculating bipartition costs. a) The internal nodes of a gene tree mapped to the internal nodes of a species tree. There is one singly mapped and one multiply mapped species tree node. The multiply mapped node must be locally reconciled, but the `BipartitionCost` algorithm (Algorithm 2) needs to choose which of the two discordant branches of the gene tree to reconcile first (labelled with a red “1” and “2”). b) Calculating the `BipartitionCost` of edge 1. Choosing this edge will produce two NNI-rearranged gene trees, shown here (along with the bipartition plane shown as a red vertical line). Under each of the two trees are shown the four scores/costs that must be calculated: `Left_bipartition`, `Right_bipartition`, `Left_lost`, and `Right_lost`. Bipartition scores represent the number of bipartitions in that set that are not present in the species tree below the node being reconciled. Lost costs represent the number of losses in that set compared to the species tree below the node being reconciled. The final bipartition cost for each is simply the sum of the four costs. c) Calculating the `BipartitionCost` of edge 2. Since the lowest cost is one of the trees associated with edge 2, we would locally reconcile this edge first.

#### Choosing among events

In order for our algorithm to more efficiently and optimally choose among different evolutionary events (e.g. Figure 2b vs 2c in the main text), we introduce the concept of a “local reconciliation” for a multiply mapped node. Here, a species tree node is locally reconciled if the number of gene tree nodes mapped to it, *m*, is reduced. For internal nodes of the species tree, this, by definition, also reduces the number of discordant branches in the gene tree.

The use of local reconciliation has several advantages. Most importantly, it provides a criterion for moving forward: we can determine if a reconciliation has been achieved within a single round of the algorithm. It also makes it clear that there are some scenarios that can only be reconciled using duplication and not NNI (e.g. a multiply mapped tip node), in which case choosing events is straightforward. In a single round of our algorithm, we attempt to locally reconcile a single multiply mapped node using both NNI+loss and duplication+loss (Figure 2).

Note that, given this definition, a species tree node can be locally reconciled by NNI while still remaining multiply mapped; e.g., if *m*= 3 is reduced to *m*= 2, the node still requires more reconciliation. In this case our algorithm will carry out up to *k* = (*m* - 1) NNI moves within a single round, as long as each one results in a local reconciliation; intuitively, the number of discordant branches (*k*) should be resolvable by at most this number of NNIs. For example, starting with *m*= 3, the first NNI move will locally reconcile a node if one of the two NNI topologies reduces the number of mappings to the species tree (we have never observed a case where both NNI topologies reduce this number). In this case, *m*= 2, and we now carry out another NNI move on the updated gene tree. Only one move needs to be examined at this point, however, since the back-NNI rearrangement would return us to the starting state. The cost of all of these NNI moves is then compared to the cost of duplication and loss at the end of the round. Duplication+loss naturally allows for more than one of each type of event for a local reconciliation, and is therefore not handled differently when *m*> 2.

Finally, at the end of a round, `reconcILS` will sum the cost of all events to choose the optimal local reconciliation. If there are equally optimal reconciliations, in order to be conservative (i.e. more similar to DL reconciliation), reconcILS will report the duplication+loss solution. In the base simulations on the 40-tip tree, 4.9% of branches had equally optimal reconciliations.

#### Labeling a reconciled gene tree

A full reconciliation of a gene tree, *G*, and species tree, *S*, will have reconciled all multiply mapped nodes, leaving no discordant branches. The total number of duplications, losses, and NNIs required for this reconciliation is the sum of each event type across all branches; these numbers are reported by `reconcILS`. We also aim to label the gene tree with these events—identifying where each event occurred in history—as this is an important output of reconciliation algorithms. As `reconcILS` traverses the species tree finding optimal local reconciliations for each branch, it also records the number and location of all events. Labeling therefore consists of placing these events on the gene tree. Traditional DL reconciliation algorithms label gene tree nodes as either *D* (for duplication) or *S* (for speciation); they may also label the location of losses, *L*. Here, we additionally aim to label the NNI events induced by ILS (*N*).

Importantly, there is no standard way to label ILS events on a tree, largely because there is no exact map between gene tree nodes and species tree nodes under ILS. A branch undergoes NNI (and ILS), not a node, so it is not clear which node flanking such a branch should be chosen to be labelled. This labeling problem therefore differs fundamentally from labeling under DL and DTL algorithms because of the lack of a one-to-one map between nodes in the two trees. Here, we have created our own labeling system for ILS, one that we think is both understandable and easy to work with.

Our solution for placing the *N* label on nodes of gene trees in `reconcILS` is to denote both a daughter branch affected and the number of NNI moves that were associated with this branch. A few examples are shown in Figure A3, which also helps to illustrate why these choices were made. Figure A3a labels the gene tree from Figure 1f—with more complete tip names—with output from `reconcILS`. As can be seen, this tree has one node (denoted with a small black dot for clarity) labeled with both duplication and NNI. Further, the labeling tells you which daughter branch was reconciled with NNI (*c_2_*), and how many NNI moves were required to reconcile it (2). Without such information, simply labeling the node with *N* would not have distinguished histories involving coalescent events along the *A* branch from those along the *c_2_* branch.

`reconcILS` also labels duplications and losses in the absence of ILS, and ILS in the absence of duplication and loss. Figure A4 shows how both individual duplications and losses are labelled: observe that neither requires a label with a specified number of events (like the *N* label), nor does either need to specify on which branch they act. `reconcILS` places the label *L* on any node in the implied gene tree that is not represented by a gene copy; for example, implied tip nodes *A_l_*, *c_l_* and *B_2_* in Figure A4b do not exist in the input gene tree itself, but are labelled with an *L*. `reconcILS` can do this because every duplication event is represented internally by generating a new copy of the species tree below the duplicated node—if losses occur on these implied trees, we can track their timing as well. Note that internal nodes can also be labelled with an *L* (Figure A4c), so that the timing of loss events must be inferred by comparing the state of parent and daughter nodes (i.e. losses on internal branches will lead the descendant internal node and all of its daughter nodes to be labelled with an *L*). Finally, any node without a label can be inferred to be a speciation node, with some caveats (see next section).

**Figure A3.**
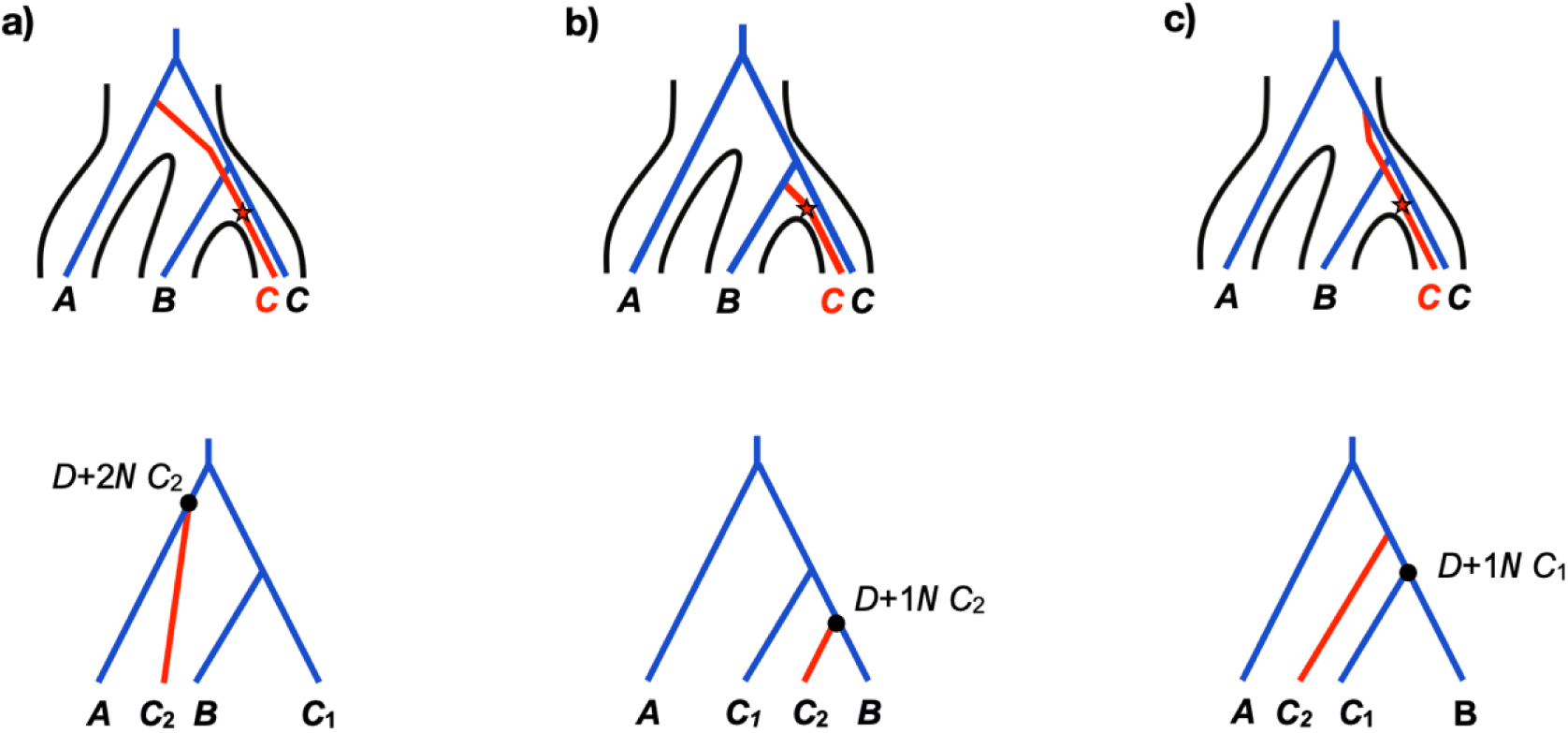
Labeling reconciled gene trees. a) The top view shows the gene tree inside the species tree (with the duplication event labeled), while the bottom view shows the reconciled gene tree labeled by `reconcILS`. The single ILS node ( ) has been labeled with a branch that has been rearranged by NNI ( ), the number of NNIs it has undergone ( ), and the fact that it also represents a duplication ( ). b) A similar example with a different gene tree history. c) Another example with a different gene tree history, though the wrong node has been labeled as the duplication node.

**Figure A4.**
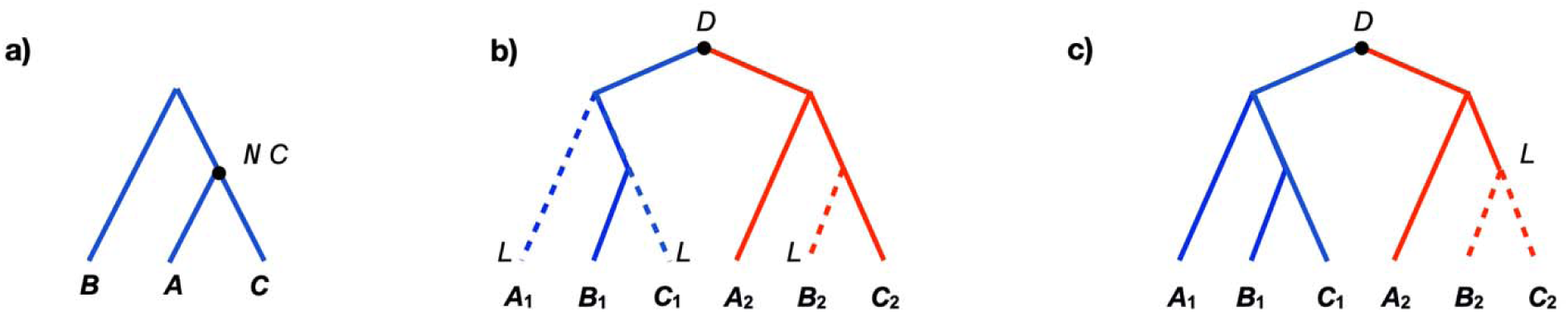
Examples of node labelling when duplication and loss do not interact with ILS. a) A labeled reconciled gene tree with only a single ILS event (and single discordant branch; as in Figure 2a in the main text). `reconcILS` chooses the node flanking the discordant branch closest to the tips to label as *N*, and further arbitrarily denotes the *C* branch as having moved. b) A reconciled tree with only duplication and loss. c) A reconciled tree with only duplication and loss, but the loss occurred on the internal branch of the gene tree.

#### Caveats on labeling events

Here, we consider two important caveats to this labeling approach, one perhaps more obvious than the other. First, a specific labeling is associated with a specific reconciliation. That is, if we had found a different optimal reconciliation for the tree in Figure A4a (for instance, one duplication and two losses), we would have had different labeled events. Therefore, a labeling is always reported in the context of a particular reconciliation. In addition, it should be noted that there may be more than one possible labeling with the same number of NNI moves, so there is some stochasticity in the exact labeling that `reconcILS` reports for histories with the same reconciliation cost.

Second, as mentioned, for some ILS events it is not always clear which of the two nodes flanking a discordant branch to label with *N*: this is why there is no exact map between gene tree node labels and species tree nodes in the presence of ILS. For a discordant branch, we have arbitrarily set `reconcILS` to always choose the node closest to the tips as *N* (e.g. Figure A4a). In some cases, this choice should not matter very much. Consider the toy reconciliation in Figure 2b from the main text—should we have placed the NNI event on node *G_l_* or on node *G_2_*? In this case the choice does not seem to imply any difference in evolutionary histories, except that unlabeled nodes may be erroneously inferred to be speciation nodes (Figure A4a shows a labeling that would result).

In contrast, consider the reconciliations shown in Figure A3b and A3c (these represent the output of `reconcILS` applied to the gene trees in Figure 1d and 1e, respectively). While we have left the daughter tree branch in red for clarity, it is important to note that these trees are identical topologically, which is why `reconcILS` has labelled them in exactly the same way. The algorithm has no way to “see” the red branches, and has consequently simply picked the bottom-most node in both cases as the one to label as having undergone NNI and duplication, even though this is not precisely correct in Figure A3c. At the moment, this is a limitation of `reconcILS` (and all methods attempting to label gene trees with ILS events), but we mention other possible solutions in the Discussion.

#### Comparison of *reconcILS* and ETE3

To test the accuracy of inferences using `reconcILS` and ETE3, the base 40-taxon species tree, in addition to 10 additional species trees with the same number of tips, were used for simulations in `dupcoal` with ILS, duplication, and loss (*λ = 0.03* and *µ* = 0.03). These parameter values were initially chosen so that the simulations produced a large number of duplications and losses to assess accuracy with, not because they are biologically motivated (alternative values were also used; see below). All simulation conditions had 1000 gene trees. The 10 additional species trees were drawn randomly using the same methods as the base tree, but with heights scaled up by drawing a random number between 3-50; the base tree was originally scaled up by 10 (see “Simulated datasets” sub-section above).

To add error, we specified three levels of gene tree errors, with a Poisson number of errors drawn for each tree. The three levels corresponded to a mean of 5, 10, or 15 errors per gene tree; all errors were modeled as an NNI rearrangement. Changes on each branch of each tree were drawn independently and sequentially, such that the same branch could be rearranged multiple times. We also carried out simulations on the base 40-tip tree with very low levels of ILS by scaling all branch lengths up by 1000; to avoid extremely large gene trees under this scenario, we also scaled the duplication and loss rates down by 1000.

The default costs (*c_D_* = 1.1, *c_L_* = 1, *c_N_* = 1) were used for all analyses of simulated data for all methods, except where noted. Note that it is the relative value of costs that matters for analysis, and the absolute values do not correspond to any biological parameter.

#### Comparison of *reconcILS* and DLCpar

We also compared the results from `reconcILS` with another reconciliation method that allows for ILS, DLCpar (Wu et al. 2014). DLCpar was written in Python 2, and—possibly because we had to run it in a virtual environment—could not run on the larger gene trees produced by the 40-tip tree. Therefore, we compared the performance of `reconcILS` and DLCpar on 1000 gene trees simulated on a 16-tip species tree (we still had to remove all gene trees with 25 or more tips, as these would not run in DLCpar). We made the rate of duplication (0.003) double the rate of loss (0.0015) to ensure that there were leaves remaining on enough gene trees. The accuracy of both methods was high for duplication, loss, and ILS (Figure A5), regardless of whether the costs used were the `reconcILS` default or the DLCpar default (e.g. for loss: `reconcILS` vs. DLCpar, *ρ*= vs. 0.82; MAE=0.037 vs. 0.042; Table A1). It is important to note that DLCpar infers the number of ILS events, rather than the number of NNI rearrangements that occur within such an event (as in `reconcILS`). To ensure that a fair comparison can be made, results reported for DLCpar in Figure A5c and Table A1 are simulated vs. inferred ILS events, rather than NNI events. In both cases the methods are doing well in their respective inferences, though ILS is overcounted by DLCpar.

One additional difference between reconciliation algorithms should be mentioned here. DLCpar does not consider losses that occur above the root of a gene tree, regardless of the species tree topology; `reconcILS` does consider these events. The difference in counting of losses among algorithms arises because `reconcILS` explicitly traverses the species tree, while DLCpar (and ETE3) does not. This enables our method to track losses that occur above the root of the input gene tree, but below the root of the input species tree. However, there may be some cases in which the behavior of DLCpar and ETE3 is preferred. For instance, since gene families must originate on some branch of the species tree (e.g. Richter et al. 2018), forcing reconciliation algorithms to infer losses on branches that pre-date this appearance would be incorrect. It is not clear which approach to counting losses one should prefer when analyzing real data.

**Figure A5.**
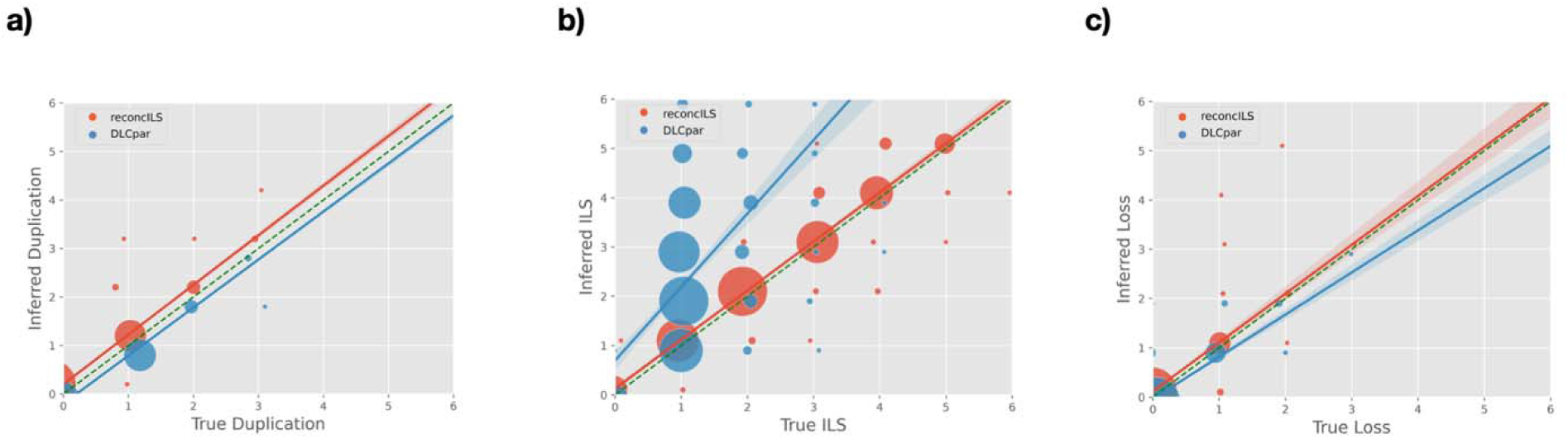
Accuracy of reconcILS and DLCpar on data simulated from the 16-tip tree. a) Number of inferred versus true duplication events. b) Number of inferred versus true ILS events. c) Number of inferred versus true loss events. Red circles represent results from `reconcILS`, blue circles represent results from DLCpar. Circle size is proportional to the number of reconciliations with each value, with jitter applied for clarity. Lines represent best-fit regressions (as calculated in Seaborn; Waskom 2021), plus bootstrap confidence intervals. The dashed line represents y=x.

**Table A1.** Comparison with DLCpar on the 16-tip tree. Each row shows results using either `reconcILS` or DLCpar for reconciliation (“method”), as well as using either the default costs from `reconcILS` or DLCpar (“cost”). The default `reconcILS` costs are *C*_D_ *= 1.1*, *C*_L_ *= 1*, and *C*_N_ *= 1*; the default costs from DLCpar are *C*_D_ *= 2*, *C*_L_ *= 2*, and *C*_N_ *= 1*. The metric “NNI” is the output from `dupcoal` counting the number of NNIs; the metric “DLCILS” counts the number of ILS events (also output from `dupcoal`), and is more similar to what DLCpar is estimating.

| #<br>Species | mu | lambda | method | cost | Metric | Spearman | MAE |
|---|---|---|---|---|---|---|---|
| 16 | 0.0015 | 0.003 | reconcILS | default reconcILS | Duplication | 0.994 | 0.015 |
| 16 | 0.0015 | 0.003 | reconcILS | default reconcILS | Loss | 0.922 | 0.037 |
| 16 | 0.0015 | 0.003 | reconcILS | default reconcILS | NNI | 0.986 | 0.089 |
| 16 | 0.0015 | 0.003 | DLCpar | default reconcILS | Duplication | 0.996 | 0.003 |
| 16 | 0.0015 | 0.003 | DLCpar | default reconcILS | Loss | 0.771 | 0.09 |
| 16 | 0.0015 | 0.003 | DLCpar | default reconcILS | DLCILS | 0.518 | 1.357 |
| 16 | 0.0015 | 0.003 | reconcILS | default DLCpar | Duplication | 0.994 | 0.015 |
| 16 | 0.0015 | 0.003 | reconcILS | default DLCpar | Loss | 0.922 | 0.037 |
| 16 | 0.0015 | 0.003 | reconcILS | default DLCpar | NNI | 0.986 | 0.089 |
| 16 | 0.0015 | 0.003 | DLCpar | default DLCpar | Duplication | 0.996 | 0.003 |
| 16 | 0.0015 | 0.003 | DLCpar | default DLCpar | Loss | 0.882 | 0.042 |
| 16 | 0.0015 | 0.003 | DLCpar | default DLCpar | DLCILS | 0.580 | 1.383 |

#### Effect of event costs on ***reconcILS***

We examined results from `reconcILS` when using different costs (the numbers in the main text all used the default costs: *C*_D_ *= 1.1*, *C*_L_ *= 1*, and *C*_N_ *= 1*). When increasing the cost of duplication and loss (*C*_D_ *= 4*, *C*_L_ *= 3*, and *C*_N_ *= 1*), the results remained largely the same, with high correlations between inferred and true duplication, loss, and NNI: *ρ*=0.93, 0.65, and 0.94, and MAE = 1.1, 1.1, 4.1 respectively. The same is true when using the default costs from DLCpar (*C*_D_ *= 2*, *C*_L_ *= 2*, and *C*_N_ *= 1*): all correlations remain high (*ρ*=0.95, 0.63, and 0.95, and MAE = 0.56, 1.36, 2.05, respectively). However, when increasing the cost of ILS (*C*_D_ *= 1*, *C*_L_ *= 1*, and *C*_N_ *= 5*), the results are much worse, closely resembling those from ETE3: *ρ*=0.78, 0.22, and 0.26, and MAE = 8.5, 16.4, 32.8, respectively. This match between `reconcILS` and ETE3 should not be surprising, as having a high cost of ILS is similar to ignoring ILS.

We assessed the sensitivity of results to the choice of costs in a second way. Using the default costs in `reconcILS`, we simulated data with equal rates of duplication and loss (*λ = 0.03* and *µ = 0.03*, respectively), unequal rates (both *λ= 0.03*, *µ = 0.01* and *λ = 0.01*, *µ = 0.03*), and highly unequal rates (both *λ= 0.03*, *µ = 0.0001* and *λ = 0.0001*, *µ = 0.03*). As can be seen in Table A2, the accuracy of `reconcILS` remains high, even when the default costs do not match the relative event frequency.

Despite this apparent robustness, alternative approaches to determining duplication and loss costs may be desired. One approach would be to infer the number of such events using an alternative, non-reconciliation approach (e.g. Mendes et al. 2020), setting the cost in `reconcILS` to be inversely related to the inferred number of events of each type. Varying rates of duplication and loss can also be inferred along different branches of the species tree, and so the costs of these events could vary commensurately along the tree. Other papers have suggested further methods for independently determining costs (e.g. Ronquist 2003; Merkle et al. 2010; Baudet et al. 2015; Sinaimeri et al. 2023).

**Table A2.** Effect of event costs on inference. The table shows results for `reconcILS` under simulations with different rates of duplication and loss. All reconciliations were carried out with default costs, even when duplication and loss rates vary by orders of magnitude. MAE: mean absolute error. Spearman: Spearman’s *ρ*.

| Simulation | lambda | mu | Event | Spearman | MAE |
|---|---|---|---|---|---|
| Dup=Loss | 0.03 | 0.03 | Duplication | 0.95 | 0.57 |
|  | 0.03 | 0.03 | Loss | 0.63 | 1.37 |
|  | 0.03 | 0.03 | NNI | 0.95 | 2.06 |
| Dup>Loss | 0.03 | 0.01 | Duplication | 0.96 | 0.66 |
|  | 0.03 | 0.01 | Loss | 0.45 | 1.72 |
|  | 0.03 | 0.01 | NNI | 0.94 | 2.25 |
| Loss>Dup | 0.01 | 0.03 | Duplication | 0.96 | 0.23 |
|  | 0.01 | 0.03 | Loss | 0.77 | 0.68 |
|  | 0.01 | 0.03 | NNI | 0.95 | 1.47 |
| Dup>>Loss | 0.03 | 0.0001 | Duplication | 0.96 | 0.60 |
|  | 0.03 | 0.0001 | Loss | 0.05 | 1.74 |
|  | 0.03 | 0.0001 | NNI | 0.94 | 2.13 |
| Loss>>Dup | 0.0001 | 0.03 | Duplication | 1.00 | 0.00 |
|  | 0.0001 | 0.03 | Loss | 0.91 | 0.31 |
|  | 0.0001 | 0.03 | NNI | 0.95 | 1.00 |

#### Runtime

Finally, we assessed the runtime and memory requirements of `reconcILS` compared to ETE3 on a single node of the Indiana University Research Desktop, running an Intel 12-core Xeon processor with 20 threads and 20 GB of RAM total (DLCpar can only be run in Python 2, which makes comparisons highly unfair). On a 16-species tree, both programs are extremely fast (average time of 0.14 seconds per gene tree for `reconcILS` and 0.22 seconds per gene tree for ETE3; Figure A6a) and both use very little memory (average peak memory per gene tree of 3.12 MB for `reconcILS` and 0.54 MB for ETE3; Figure A6b). Both programs remain very fast on data produced from a 40-species tree, though the faster speed of ETE3 is also evident (5.71 seconds per gene tree for `reconcILS` and 1.79 seconds for ETE3; Figure A6a). Regardless, this means that the entire 1000-gene-tree simulated dataset on the 40-species tree took less than 84 minutes to analyze using `reconcILS` on a moderately powerful computer. Similarly, the memory requirements of `reconcILS` increase on the 40-tip tree (33.22 MB per gene tree for `reconcILS` and 1.92 MB for ETE3; Figure A6b), but are not onerous for modern computers. As the 40-species tree can contain many more events (duplication, loss, or ILS), we also plotted the average time and memory required for trees containing different numbers of total events (Figure A6c,d). Both programs show approximately linear increases in time and memory with the number of events, which is promising for analyzing larger datasets with larger species trees, though we have not systematically tested it on much larger trees.

**Figure A6.**
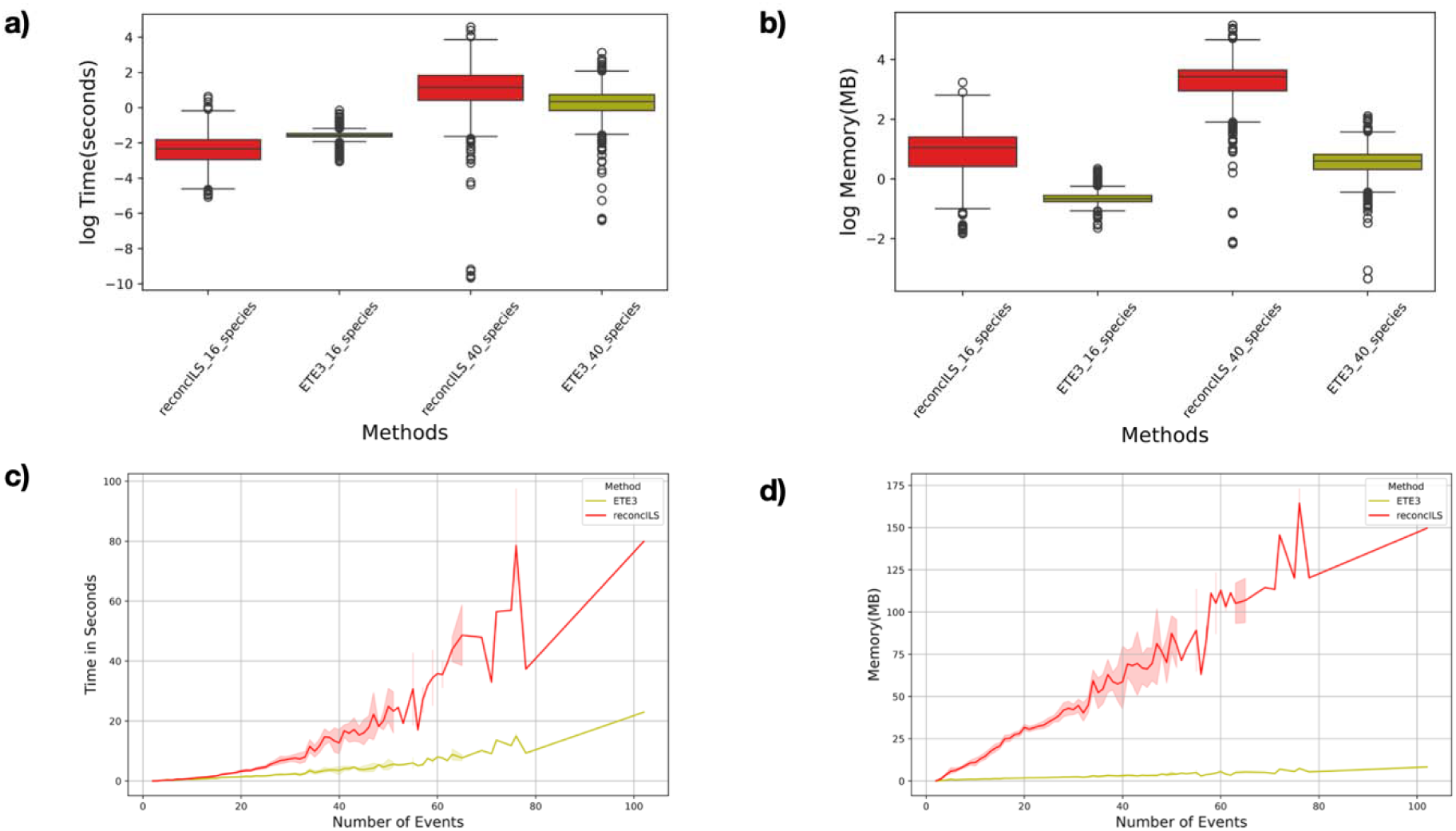
Run-time and peak memory consumption for reconcILS and ETE3. a) Comparison of the run-time to reconcile individual gene trees from species trees with either 16 species or 40 species. b) Comparison of peak memory consumption to reconcile individual gene trees from species trees with either 16 species or 40 species. c) Run-time as a function of the number of events (ILS, duplication, loss) per gene tree for only the 40-species tree. d) Peak memory consumption as a function of the number of events (ILS, duplication, loss) per gene tree for only the 40-species tree.

## Algorithms

### Notation and terminology

1. *G* and *S* denote the complete gene tree and species tree, respectively.
2. Lowercase *g* ε *V(G)* and s ε *V(S)* always denote *nodes.*
3. *G\** and *S\** denote a proposed NNI-rearranged gene subtree and its corresponding LCA-mapped species subtree.
4. The cost vector *(C_L_, C_D_, C_N_)* specifies the costs of a loss, a duplication, and one NNI operation, respectively.
5. The *bipartition score* is the number of bipartitions in a, gene tree subtree that a.re absent from the species tree. The *bipartition cost* (*σ*) is the sum of the left and right bipartition scores and their induced loss costs.
6. A *discordant branch* is flanked by gene tree nodes mapped to the same species tree node. If *m* gene tree nodes map to a particular species tree node, then *k* = *m* - 1 is the number of discordant branches that must be locally reconciled.

#### Algorithm 1

**reconcILS: Iteratively reconciles a, gene tree (*G*) with a, species tree (*S*) using duplication, loss, and NNI operations**

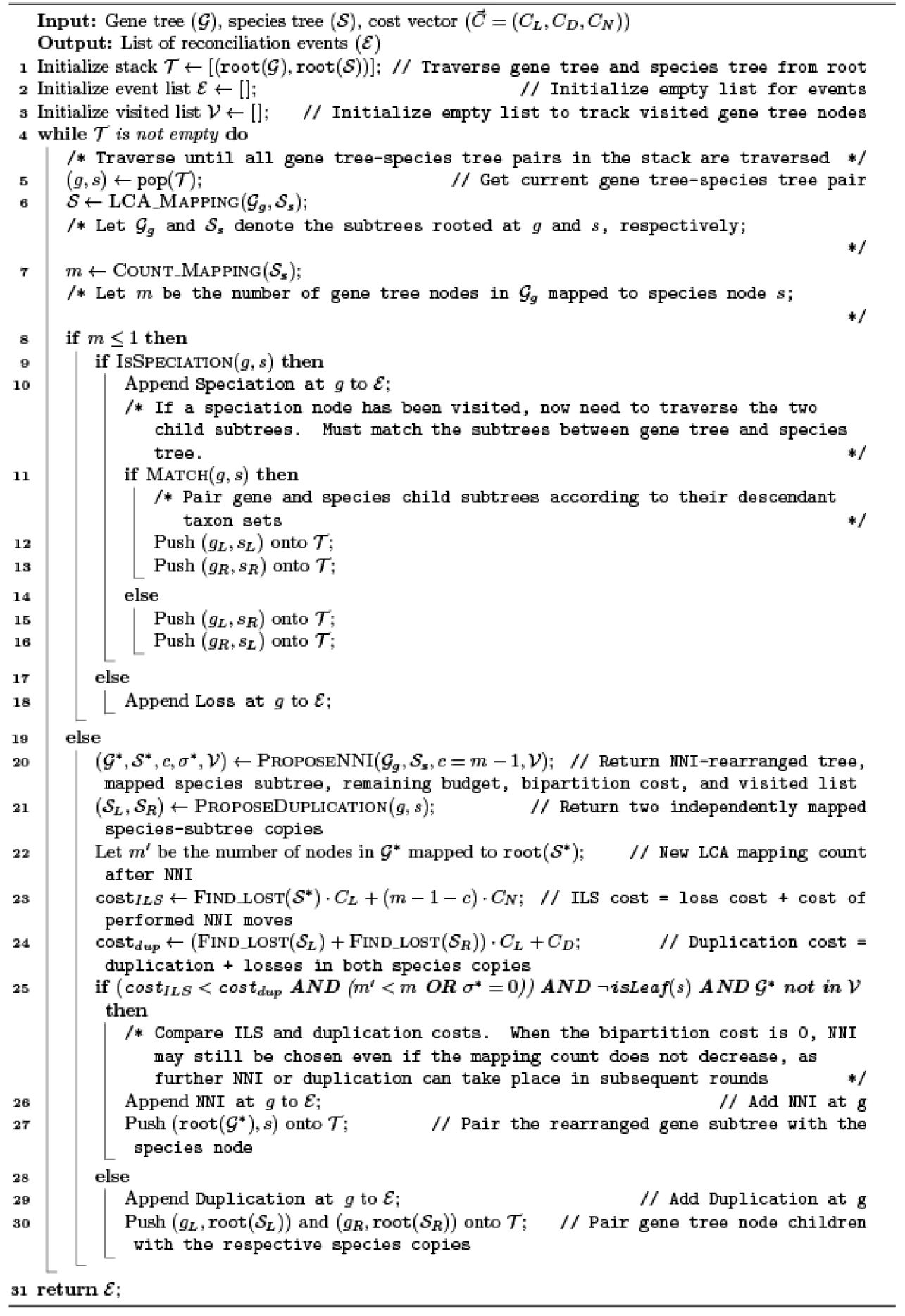

#### Algorithm 2

**BipartitionCost: Identifies discordant branch in *G* whose NNI rearrangement yields lowest bipartition cost with S* using mapping count and depth as tie-breakers**

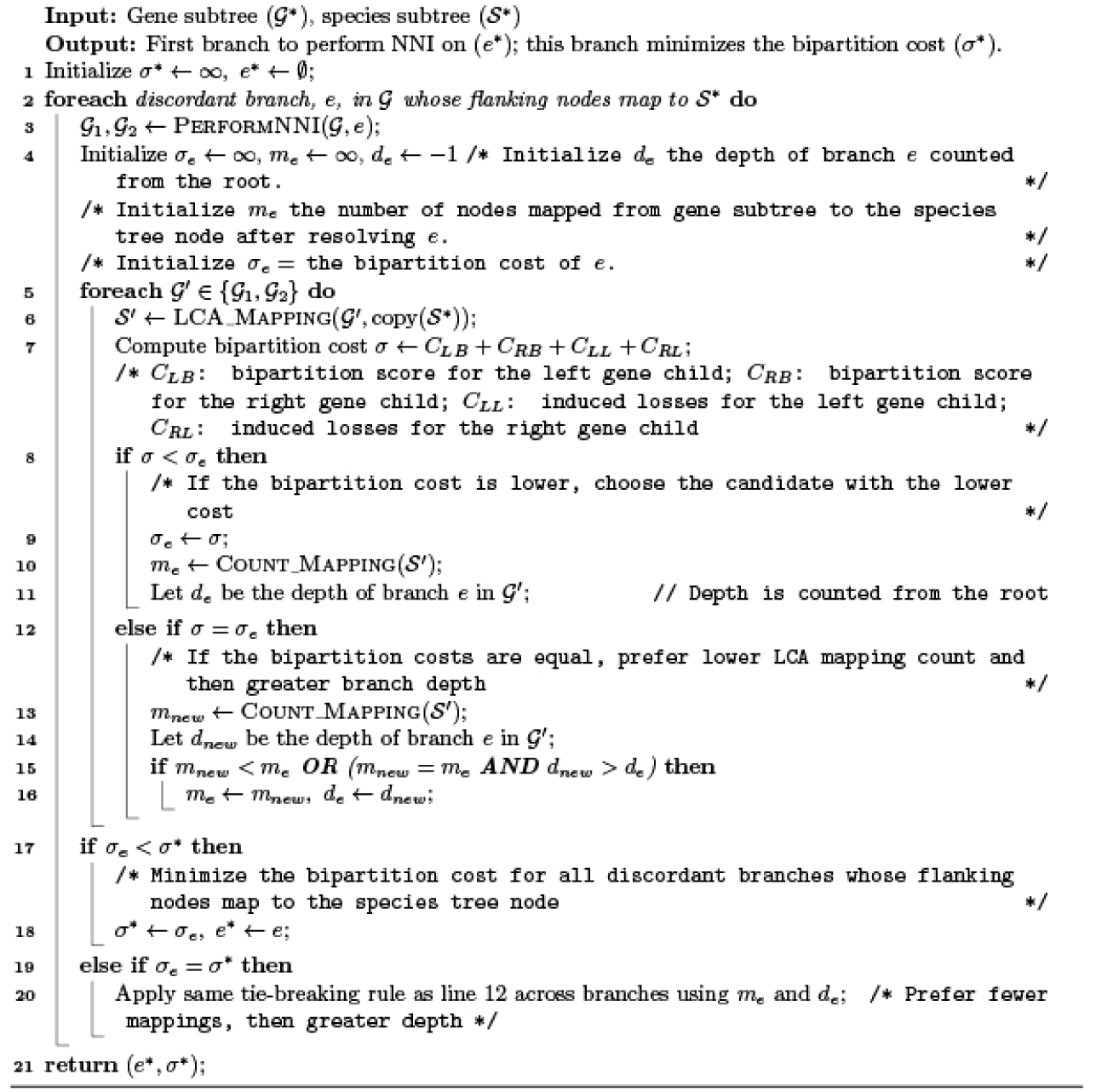

#### Algorithm 3

**ProposeDuplication**

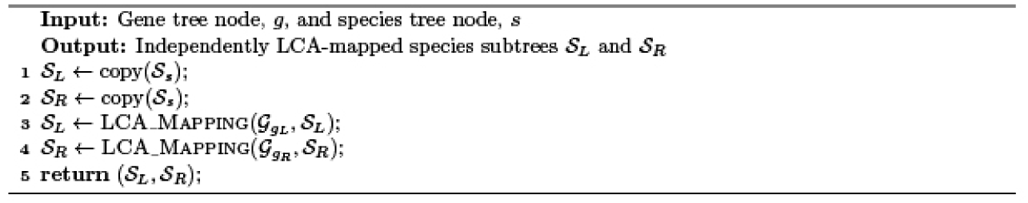

#### Algorithm 4

**ProposeNNI**

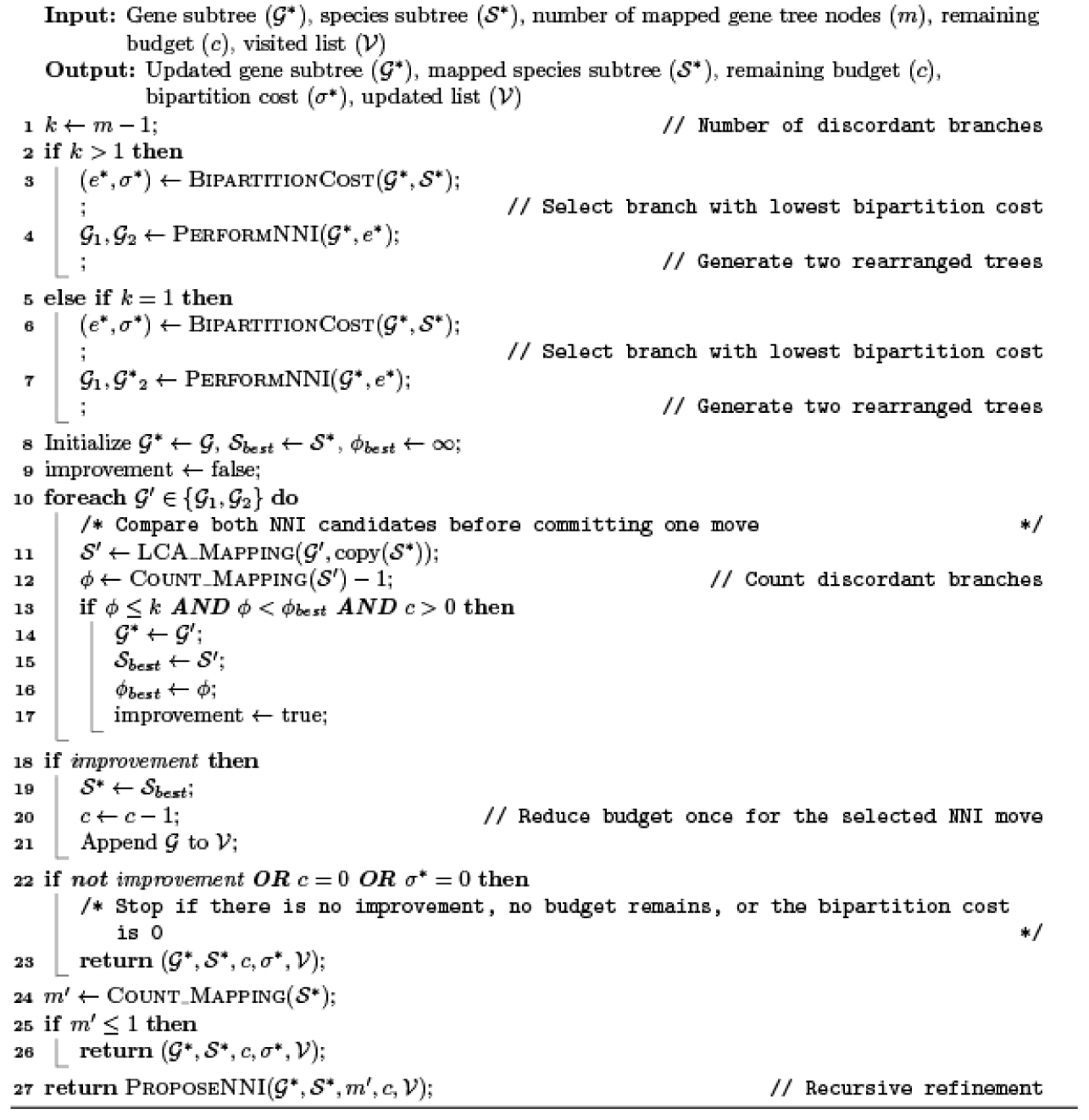

#### Algorithm 5

**PerformNNI**

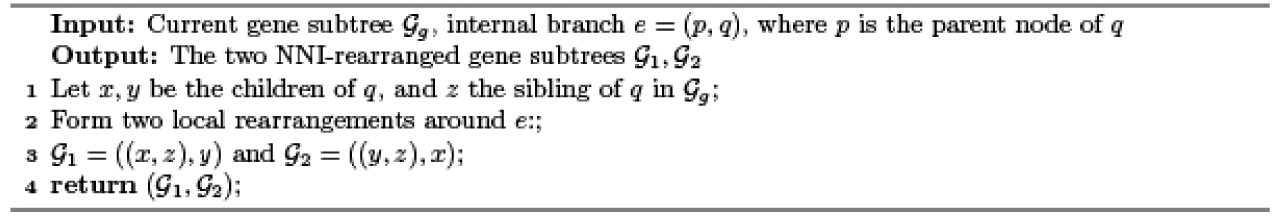

#### Algorithm 6

**Helper Functions**

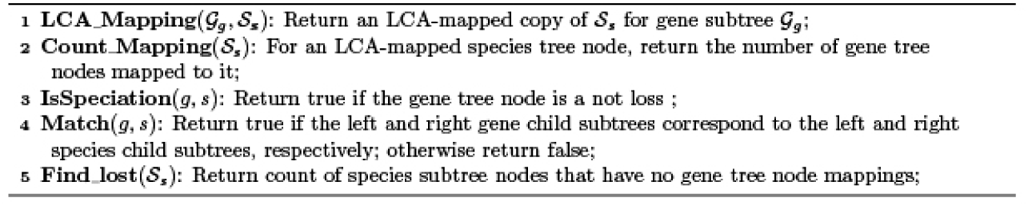

## Supplementary Materials

## Supplementary Tables

***Table S1. Adding gene tree error or removing ILS*.**

The table shows results from the standard run (no error, regular branch lengths), as well as datasets with increasing levels of error (runs with a mean of 5, 10, or 15 errors per gene tree) and a dataset in which branch lengths were scaled by 1000, so that there is almost no ILS. Results from `reconcILS` and ETE3 are shown as mean absolute error (MAE).

***Table S2. Accuracy of reconciliation on simulations using 10 additional species trees*.**

The table shows results from running `reconcILS` and ETE3 on 10 additional 40-species trees. The height of each tree (in coalescent units) is reported under “tree height”. Results are shown as both Spearman’s *ρ* and mean absolute error (MAE).

---

**Figure S1.**
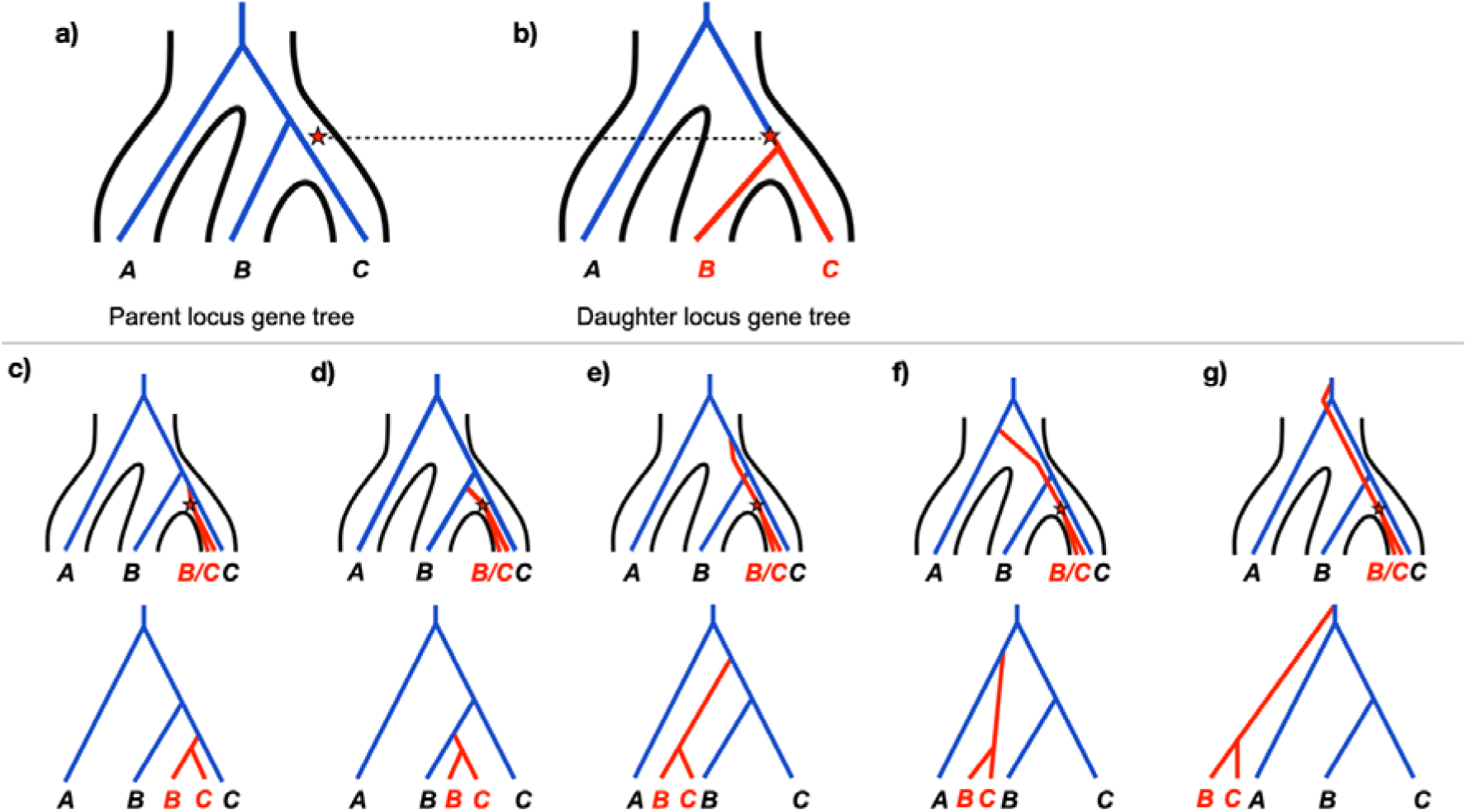
The MSC-DL model generates full gene trees. This figure mirrors Figure 1 in the main text, but the duplication inserts on an internal branch of the gene tree at the daughter locus. a) A gene tree at the parent locus (blue) is drawn from the species tree (outlined in black) via the multispecies coalescent (MSC) model. A duplication event (red star) is drawn from a birth-death model, in this case occurring in the population ancestral to species and . b) A gene tree at the daughter locus is drawn from the MSC. The duplication is placed on this gene tree on the branch and at the time at which it occurred in the species tree (shown as a dotted line). The inserted DNA now takes on the gene tree branch labeled in red. c)-g) Possible ways that the parent and daughter loci coalesce to form the full gene tree. In all panels the star and red branch below it come from the daughter gene tree, and the blue tree comes from the parent gene tree. The coalescent process joining the two trees is shown as a red branch above the star. The trees in the bottom row are simply unfolded views of the full gene trees.

**Figure S2.**
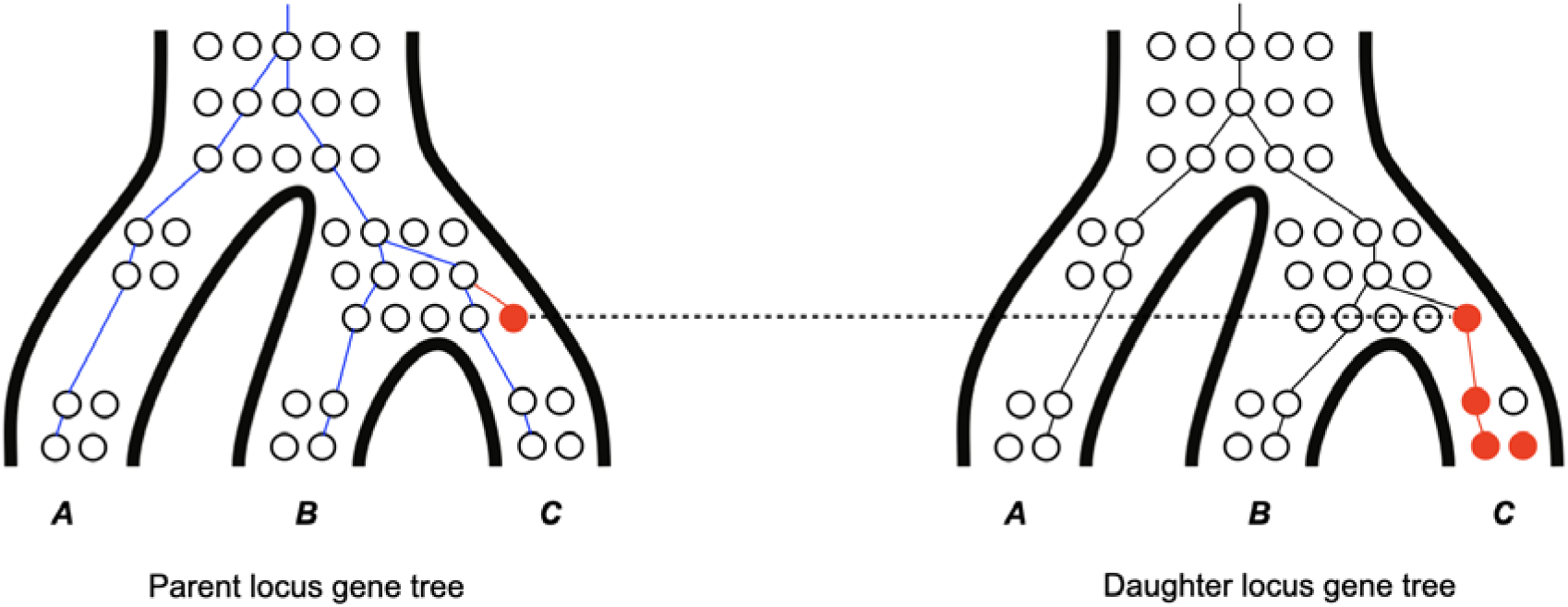
Population genetic view of the MSC-DL model. A population genetic view of the scenario in Figure 1c from the main text. At the parent locus, the blue lines describe the sampled gene tree. Individuals are represented as circles. One individual is chosen as the donor of the duplication (filled red circle), and the red line describes the coalescence of that individual with the others sampled at the parent locus. At the daughter locus, the duplication is inherited by the individual in which the mutation occurred (denoted by the dashed horizontal line). The duplication can then rise in frequency in subsequent generations, with the red line connecting those individuals included in the sampled gene tree at this locus. The full gene tree corresponds to the red and blue gene trees together. Sampled individuals at the daughter locus without the duplication have a genealogy (in black), but this is not relevant for the full gene tree.

**Figure S3.**
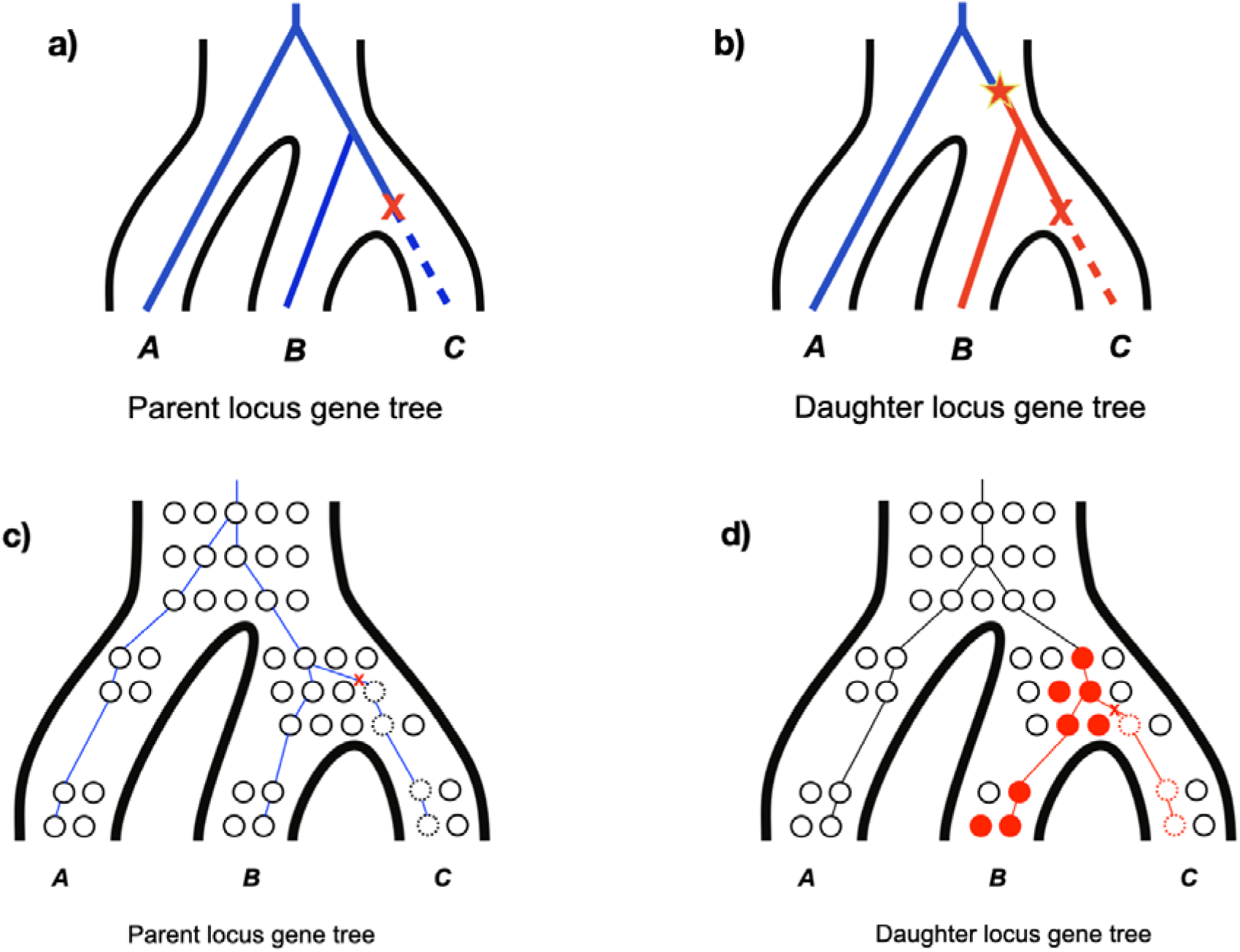
Loss events. a) Loss events occurring at the parent locus. b) Loss events occurring at a daughter locus. Note that a duplication event must precede a loss at a daughter locus in order to be observed. c) and d) Population genetic view of panels a) and b), respectively. Dotted lines around individuals (circles) indicate that those individuals no longer carry genetic material at the locus. In panel d), filled circles indicate individuals that have inherited the duplicative mutation.

**Figure S4.**
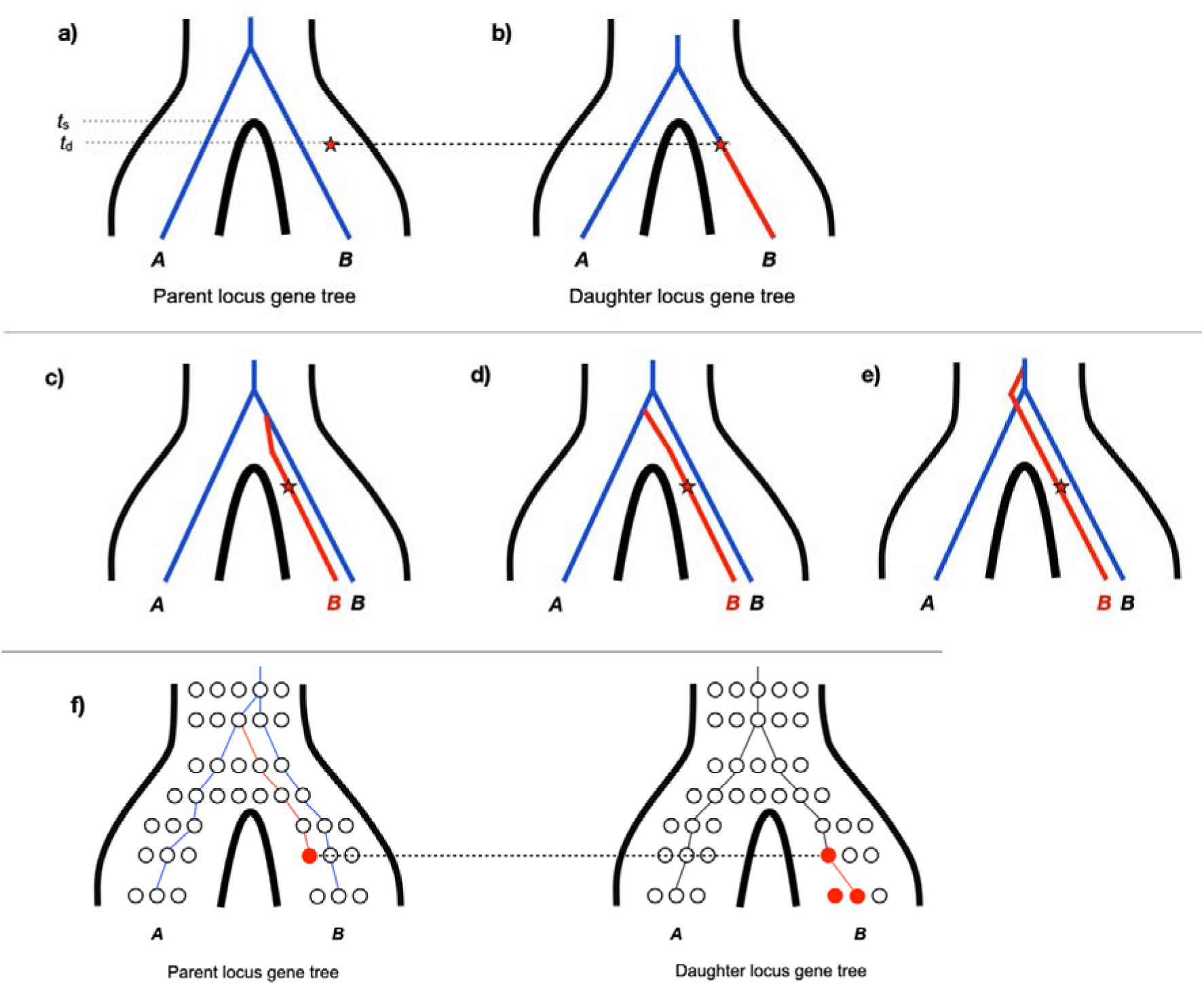
Unsorted homology in two species (with ILS). a) A duplication occurs at the parent locus and inserts b) at the daughter locus. The time of speciation (*t*_s_) and the time of duplication (*t*_d_) are also shown. c-e) Three ways that the duplicate can coalesce with the gene tree at the parent locus. In all three panels there is ILS, since the duplicate and the parent do not coalesce while in the *B* tip branch of the species tree. In panels d) and e) there is also unsorted homology, as the duplicate in species *B* does not coalesce with the gene tree *B* branch at the parent locus. f) Population genetic view of the scenario depicted in panel d). See Figures S2 and S3 for an explanation of symbols.

**Figure S5.**
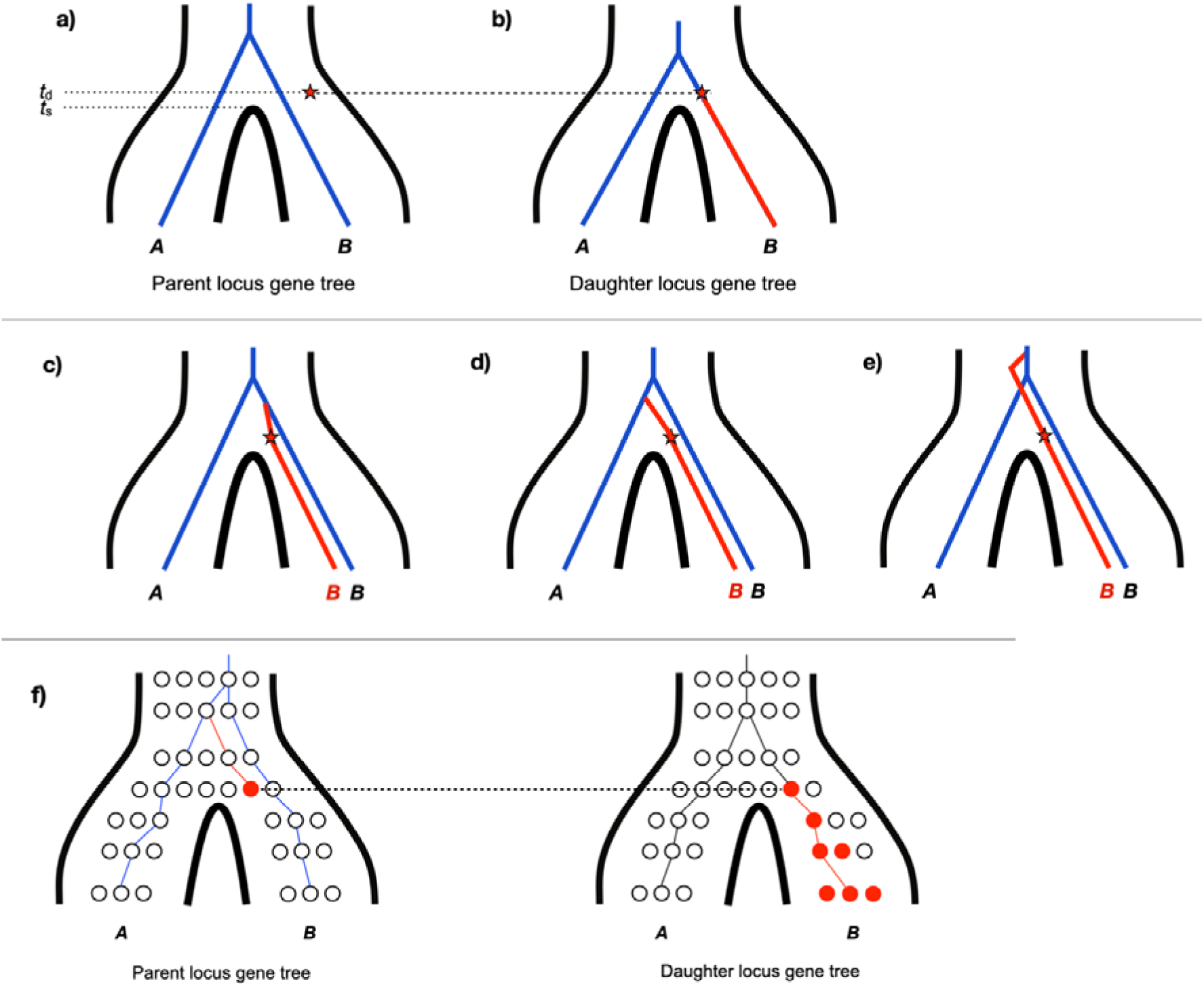
Unsorted homology in two species (no ILS). a) A duplication occurs at the parent locus and inserts b) at the daughter locus. The time of speciation (*t*_s_) and the time of duplication (*t*_d_) are also shown. c-e) Three ways that the duplicate can coalesce with the gene tree at the parent locus. In all three panels there is no ILS, since the duplicate and the parent coalesce in the shared *AB* population in which the duplication occurred. In panels d) and e) there is also unsorted homology, as the duplicate in species *B* does not coalesce with the gene tree *B* branch at the parent locus. f) Population genetic view of the scenario depicted in panel d). See Figure S2 and S3 for an explanation of symbols.

**Figure S6.**
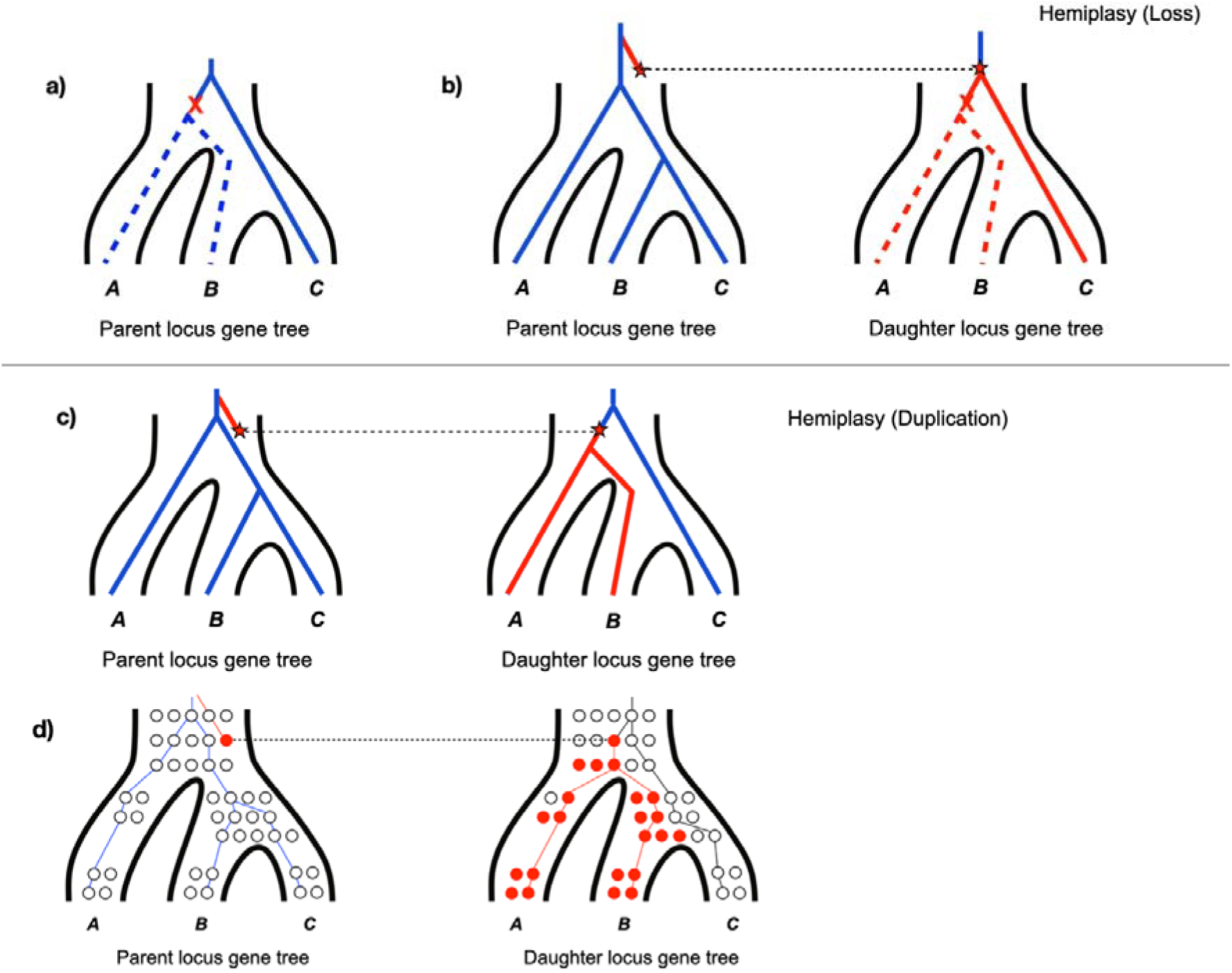
Copy-number hemiplasy for duplication and loss. Copy-number hemiplasy (CNH) occurs when a mutation lands on a discordant branch of a gene tree. For losses, CNH can occur at a) the parent locus, or b) the daughter locus (following a duplication). For duplications, CNH can only occur at c) the daughter locus. d) Population genetic view of the scenario depicted in panel c). See Figure S2 and S3 for an explanation of symbols.

**Figure S7.**
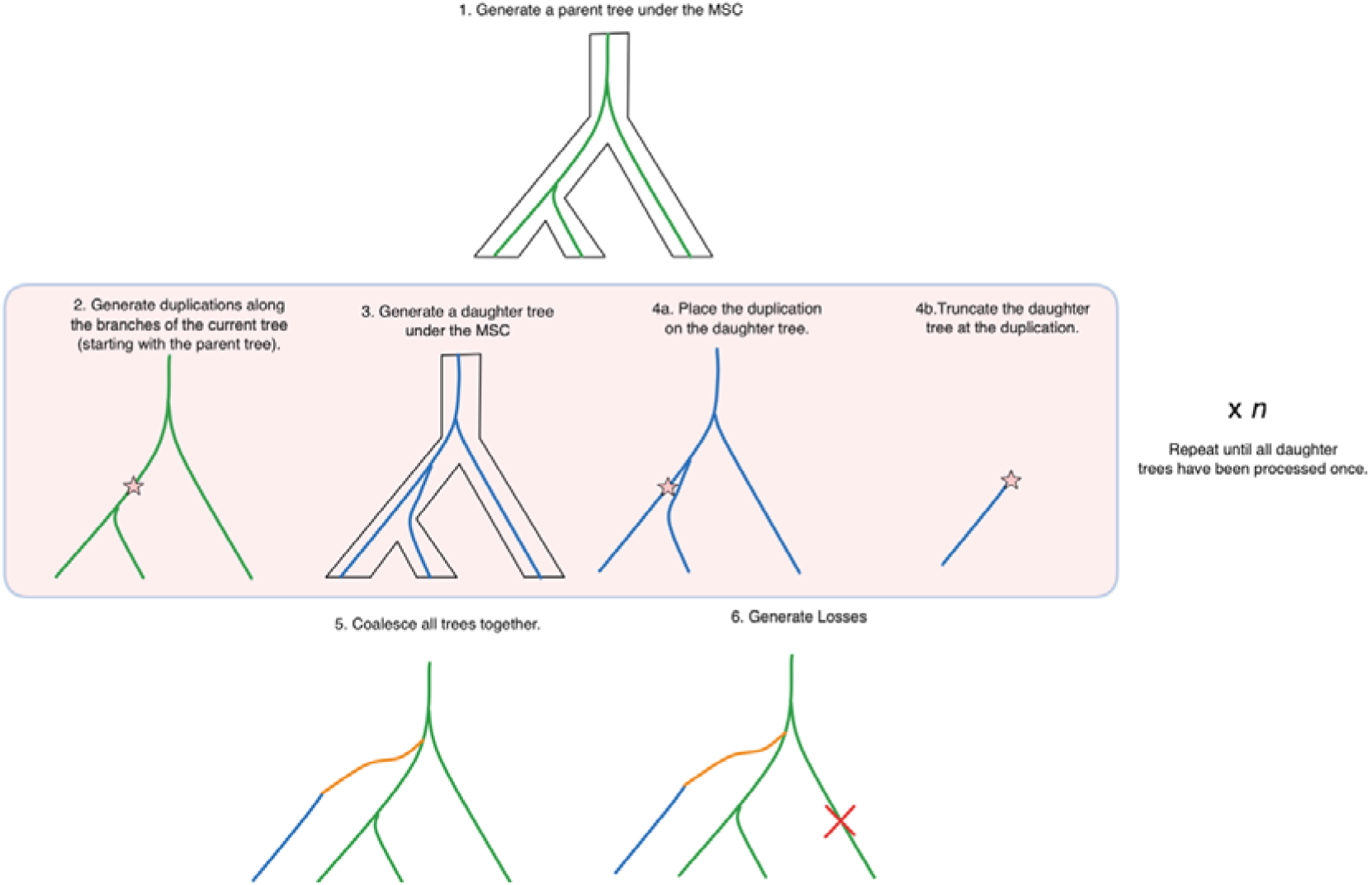
Overview of dupcoal simulation process. Details on the `dupcoal` algorithm can be found in the Appendix. Here, we give a pictorial overview of the simulation steps.

**Figure S8.**
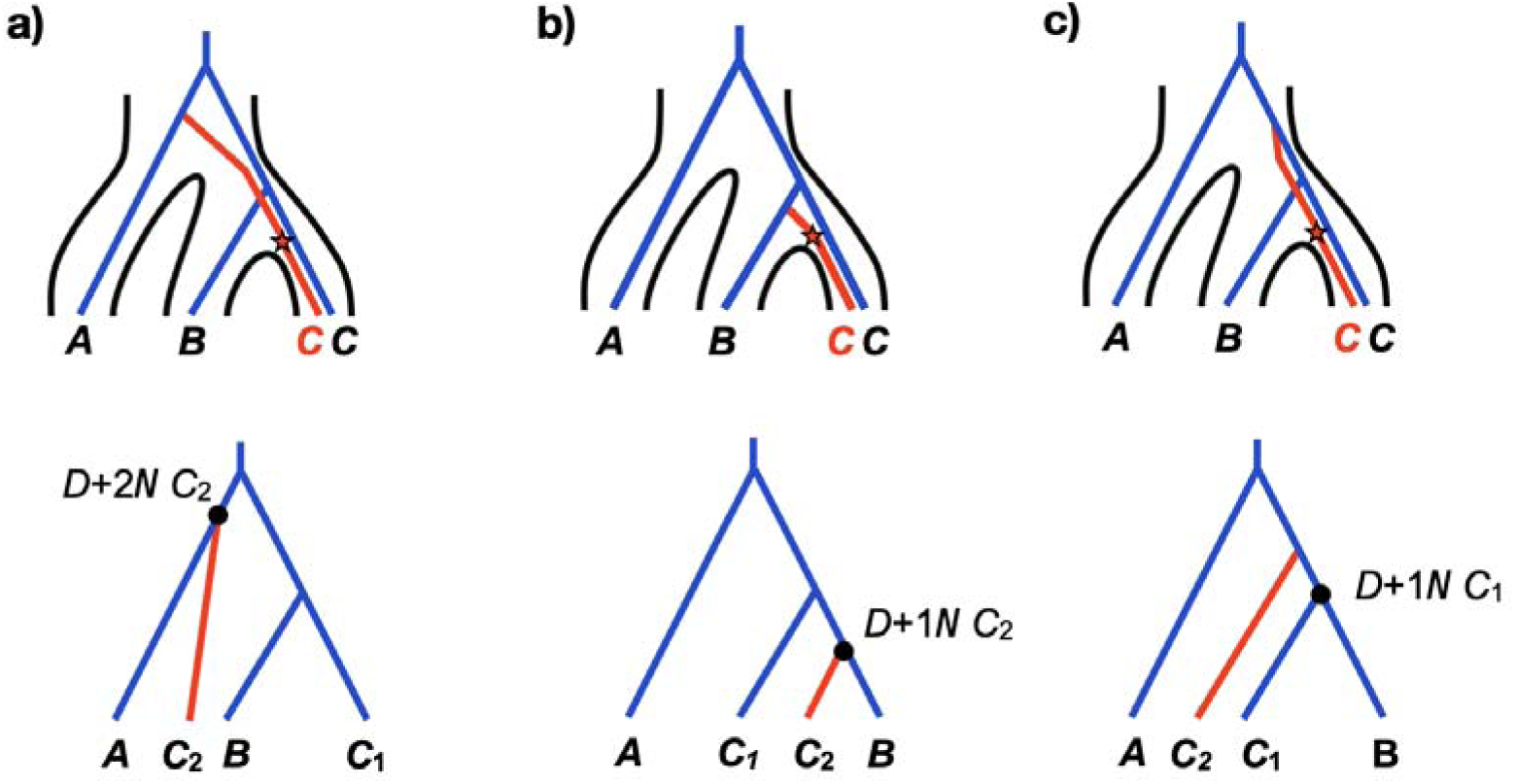
Labeling reconciled gene trees. All panels show the generation process of a gene tree on the top (similar to Figure 1 in the main text) and the labeling of the resulting gene tree by `reconcILS` on the bottom. a) One possible reconciliation of a gene tree labeled by `reconcILS`. The single ILS node ( ) has been labeled with a branch that has been rearranged by NNI ( ), the number of NNIs it has undergone ( ), and the fact that it also represents a duplication ( ). b) Similar to panel a, but with a different gene tree. This tree is correctly labeled, but note that it is the same topology as in panel c. c) Similar to panels a and b, but with a different gene tree. Note that the program has labeled the bottom-most node flanking the discordant branch with and, although that is not the true history.

**Figure S9.**
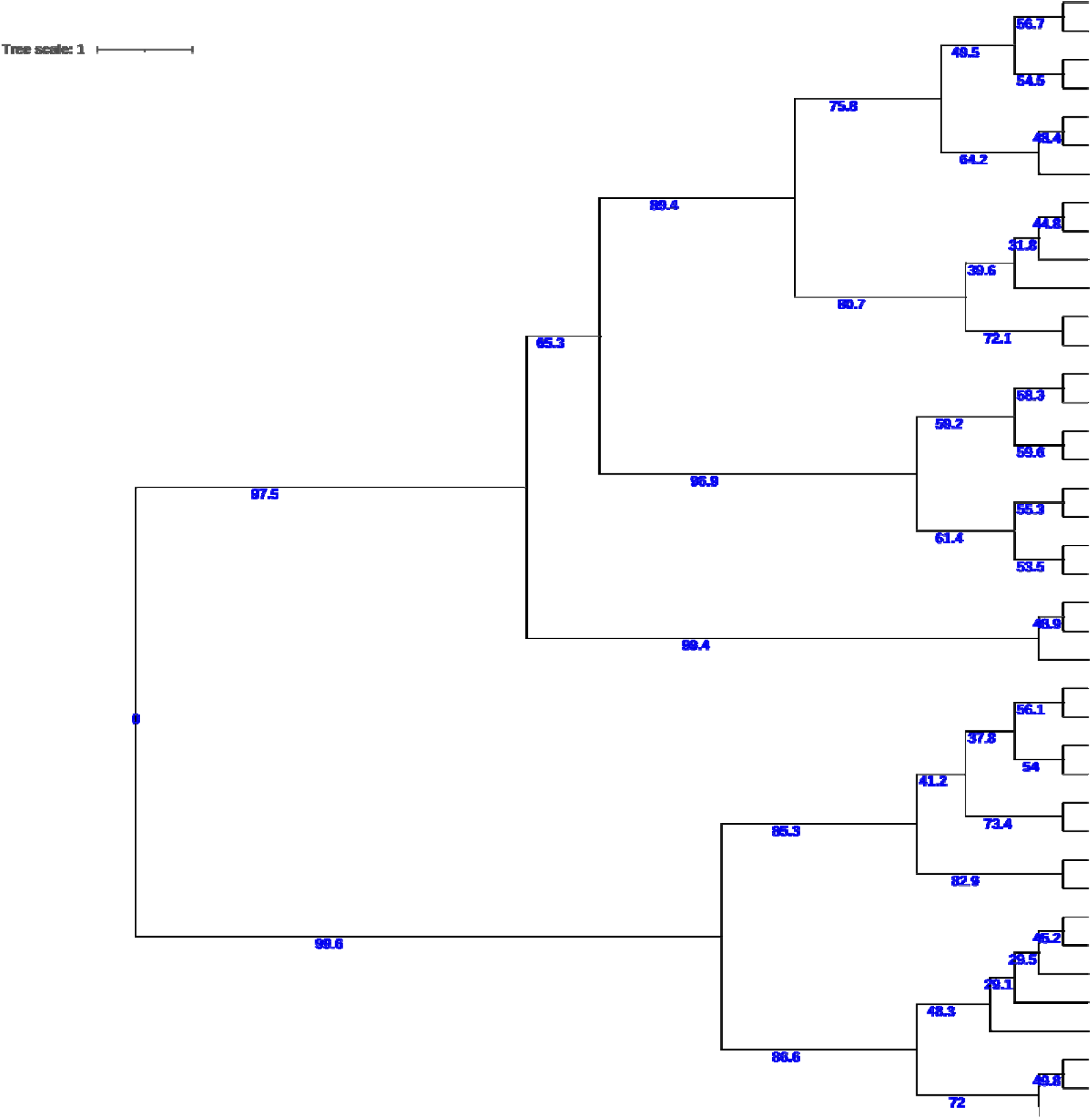
40-species tree. The main species tree used in simulation of gene trees. All branches are shown in coalescent units, with estimated concordance factors given on all internal branches.

**Figure S10.**
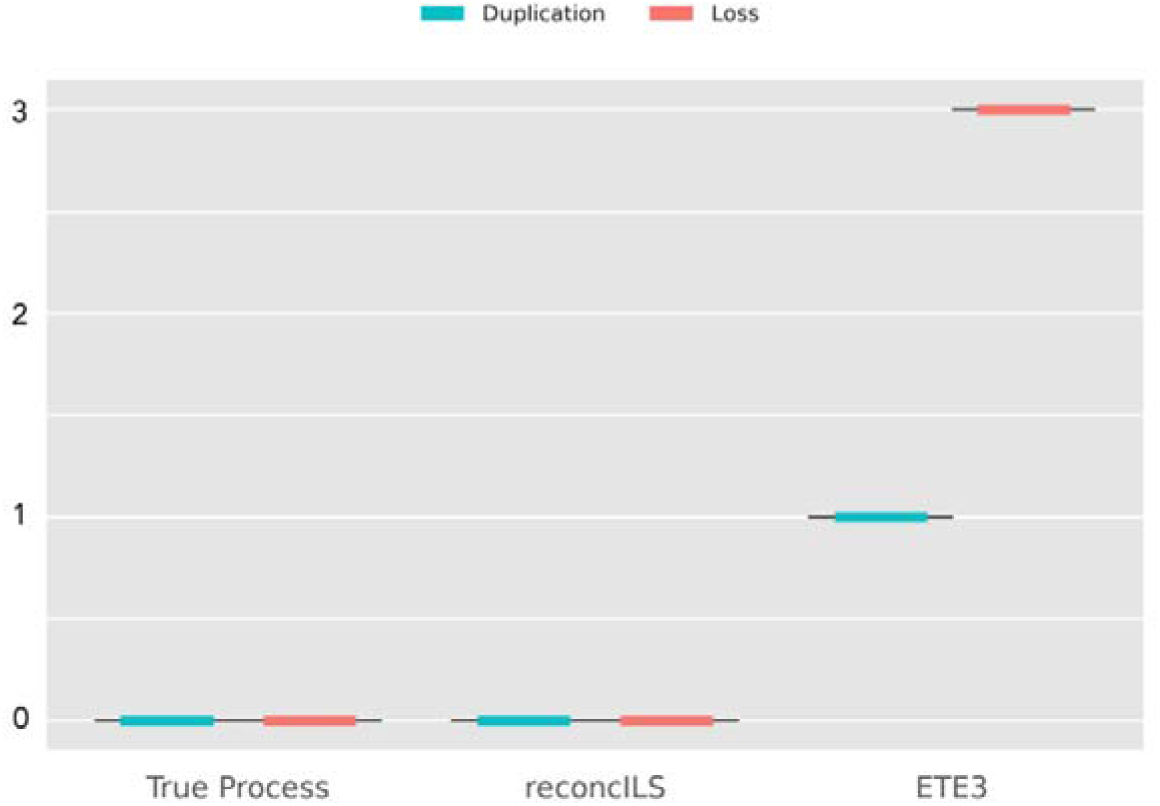
Accuracy of ETE3 on data simulated from the 3-tip tree with no duplication and loss. On 3-tip trees that are discordant solely due to ILS, `reconcILS` correctly infers no duplications or losses. In contrast, ETE3 (and all DL-reconciliation methods) will infer one duplication and three losses. Results are from 1000 gene trees, but there is no variance in outcomes.

**Figure S11.**
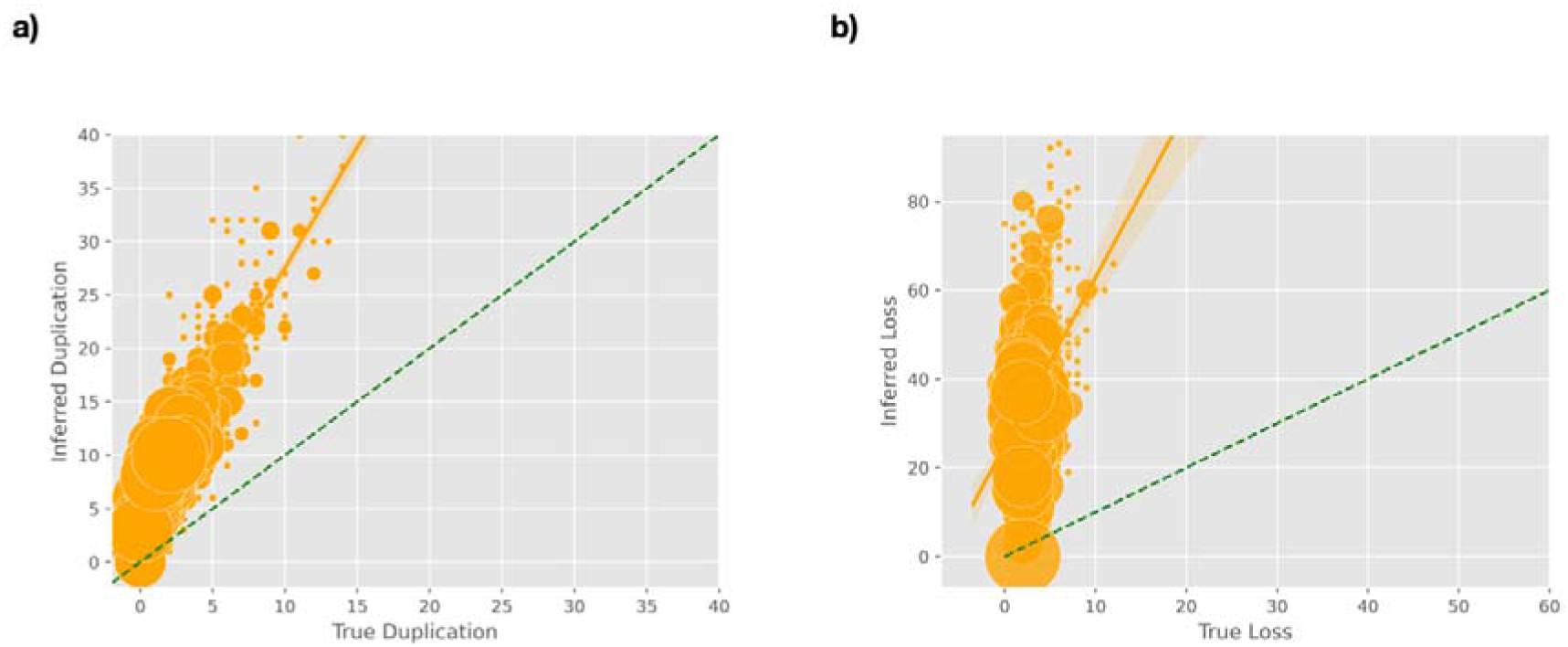
Accuracy of ETE3 on data simulated from the 40-tip tree. a) Number of inferred versus true duplication events. b) Number of inferred versus true loss events. Circle size is proportional to the number of reconciliations with each value, with jitter applied for clarity. The yellow lines represent best-fit regression (as calculated in Seaborn; Waskom 2021), plus bootstrap confidence intervals. The dashed lines represent y=x.

**Figure S12.**
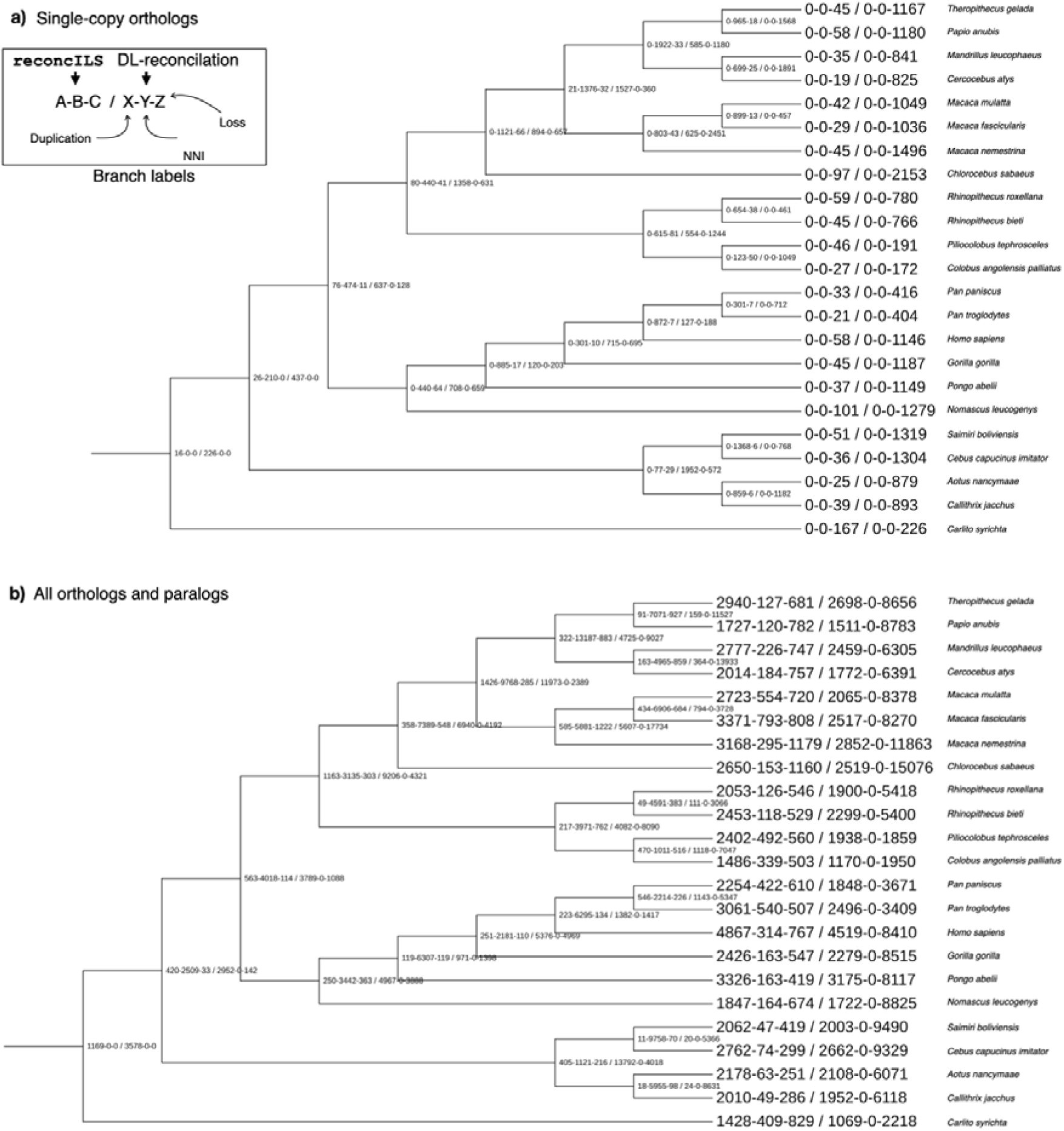
Numerical results from primates. a) Analysis of 1,820 single-copy orthologs. Both `reconcILS` and ETE3 (a DL-reconciliation method) were used to analyze the data, with duplication, NNI, and loss events mapped to each branch of the species tree. On each branch, there are two sets of results, each with three numbers (see inset). The first three values come from `reconcILS`, while the second three come from ETE3. Duplication, NNI, and loss are shown for both, though all NNI values for ETE3 must be 0. Values for each branch are shown to its right. b) Analysis of 11,555 gene trees containing both orthologs and paralogs. Results are presented in the same manner as in panel a.

**Figure S13.**
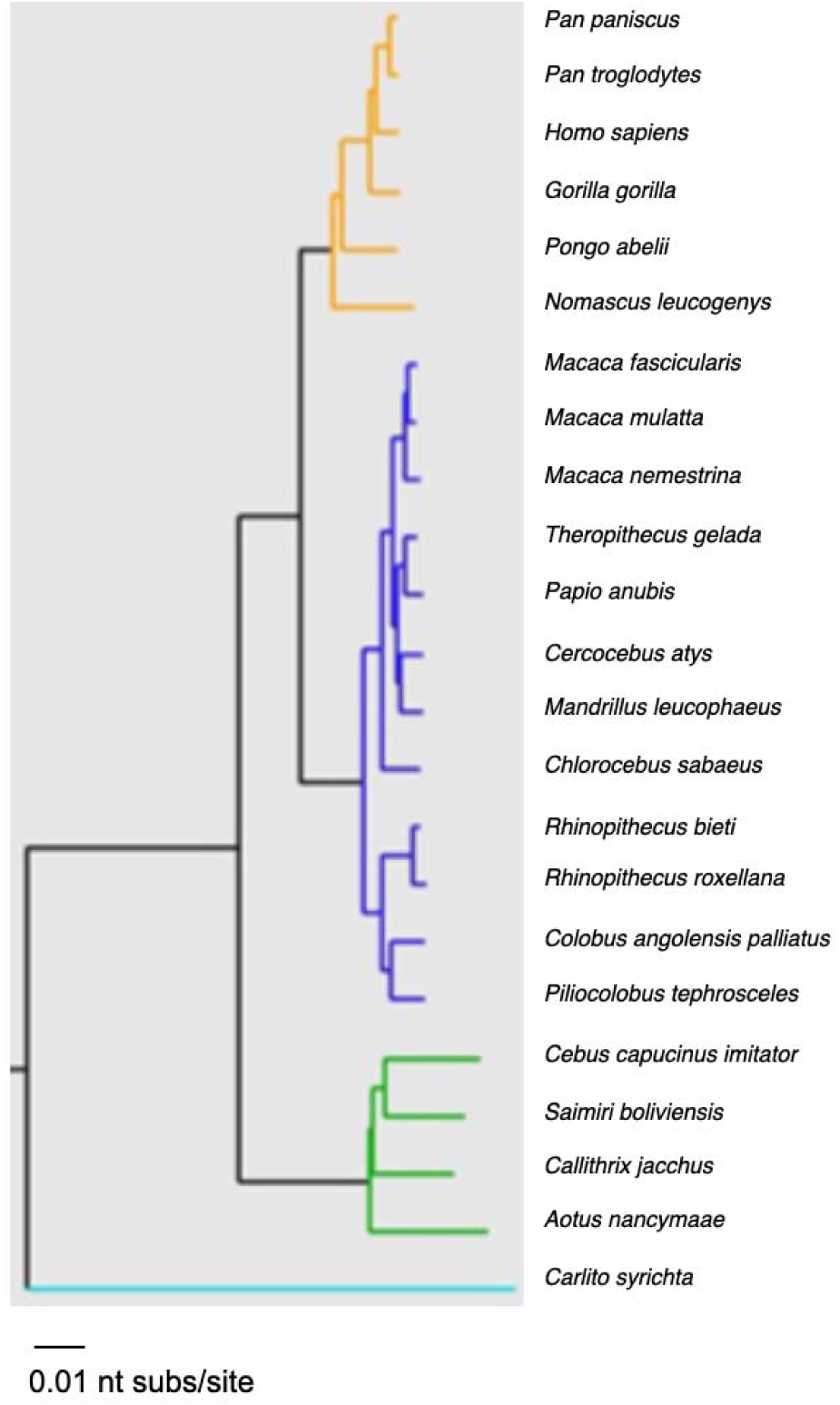
Primate tree. This the same tree as in Figure 4 and Figure S12, but with branch lengths shown in substitutions per site. From Vanderpool et al. (2020).

