## Supplementary material for "reconcILS: A gene tree-species tree reconciliation algorithm that allows for incomplete lineage sorting": Table S2

| dataset | tree height | reconciliati | simulation | mu | lambda |
|---|---|---|---|---|---|
| 1 | 45 | reconcILS | dupcoal | dupcoal: 0.03 | dupcoal: 0.03 |
| | | reconcILS | dupcoal | dupcoal: 0.03 | dupcoal: 0.03 |
| | | reconcILS | dupcoal | dupcoal: 0.03 | dupcoal: 0.03 |
| | | ETE3 | dupcoal | dupcoal: 0.03 | dupcoal: 0.03 |
| | | ETE3 | dupcoal | dupcoal: 0.03 | dupcoal: 0.03 |
| 2 | 4 | reconcILS | dupcoal | dupcoal: 0.03 | dupcoal: 0.03 |
| | | reconcILS | dupcoal | dupcoal: 0.03 | dupcoal: 0.03 |
| | | reconcILS | dupcoal | dupcoal: 0.03 | dupcoal: 0.03 |
| | | ETE3 | dupcoal | dupcoal: 0.03 | dupcoal: 0.03 |
| | | ETE3 | dupcoal | dupcoal: 0.03 | dupcoal: 0.03 |
| 3 | 20 | reconcILS | dupcoal | dupcoal: 0.03 | dupcoal: 0.03 |
| | | reconcILS | dupcoal | dupcoal: 0.03 | dupcoal: 0.03 |
| | | reconcILS | dupcoal | dupcoal: 0.03 | dupcoal: 0.03 |
| | | ETE3 | dupcoal | dupcoal: 0.03 | dupcoal: 0.03 |
| | | ETE3 | dupcoal | dupcoal: 0.03 | dupcoal: 0.03 |
| 4 | 15 | reconcILS | dupcoal | dupcoal: 0.03 | dupcoal: 0.03 |
| | | reconcILS | dupcoal | dupcoal: 0.03 | dupcoal: 0.03 |
| | | reconcILS | dupcoal | dupcoal: 0.03 | dupcoal: 0.03 |
| | | ETE3 | dupcoal | dupcoal: 0.03 | dupcoal: 0.03 |
| | | ETE3 | dupcoal | dupcoal: 0.03 | dupcoal: 0.03 |
| 5 | 16 | reconcILS | dupcoal | dupcoal: 0.03 | dupcoal: 0.03 |
| | | reconcILS | dupcoal | dupcoal: 0.03 | dupcoal: 0.03 |
| | | reconcILS | dupcoal | dupcoal: 0.03 | dupcoal: 0.03 |
| | | ETE3 | dupcoal | dupcoal: 0.03 | dupcoal: 0.03 |
| | | ETE3 | dupcoal | dupcoal: 0.03 | dupcoal: 0.03 |
| 6 | 23 | reconcILS | dupcoal | dupcoal: 0.03 | dupcoal: 0.03 |
| | | reconcILS | dupcoal | dupcoal: 0.03 | dupcoal: 0.03 |
| | | reconcILS | dupcoal | dupcoal: 0.03 | dupcoal: 0.03 |
| | | ETE3 | dupcoal | dupcoal: 0.03 | dupcoal: 0.03 |
| | | ETE3 | dupcoal | dupcoal: 0.03 | dupcoal: 0.03 |
| 7 | 10 | reconcILS | dupcoal | dupcoal: 0.03 | dupcoal: 0.03 |
| | | reconcILS | dupcoal | dupcoal: 0.03 | dupcoal: 0.03 |
| | | reconcILS | dupcoal | dupcoal: 0.03 | dupcoal: 0.03 |
| | | ETE3 | dupcoal | dupcoal: 0.03 | dupcoal: 0.03 |
| | | ETE3 | dupcoal | dupcoal: 0.03 | dupcoal: 0.03 |
| 8 | 48 | reconcILS | dupcoal | dupcoal: 0.03 | dupcoal: 0.03 |
| | | reconcILS | dupcoal | dupcoal: 0.03 | dupcoal: 0.03 |
| | | reconcILS | dupcoal | dupcoal: 0.03 | dupcoal: 0.03 |
| | | ETE3 | dupcoal | dupcoal: 0.03 | dupcoal: 0.03 |
| | | ETE3 | dupcoal | dupcoal: 0.03 | dupcoal: 0.03 |
| 9 | 46 | reconcILS | dupcoal | dupcoal: 0.03 | dupcoal: 0.03 |
| | | reconcILS | dupcoal | dupcoal: 0.03 | dupcoal: 0.03 |

| | | | | | |
|---|---|---|---|---|---|
| | | reconcILS | dupcoal | dupcoal: 0.03 | dupcoal: 0.03 |
| | | ETE3 | dupcoal | dupcoal: 0.03 | dupcoal: 0.03 |
| | | ETE3 | dupcoal | dupcoal: 0.03 | dupcoal: 0.03 |
| 10 | 3 | reconcILS | dupcoal | dupcoal: 0.03 | dupcoal: 0.03 |
| | | reconcILS | dupcoal | dupcoal: 0.03 | dupcoal: 0.03 |
| | | reconcILS | dupcoal | dupcoal: 0.03 | dupcoal: 0.03 |
| | | ETE3 | dupcoal | dupcoal: 0.03 | dupcoal: 0.03 |
| | | ETE3 | dupcoal | dupcoal: 0.03 | dupcoal: 0.03 |

| cost | Metric | Spearman | MAE |
|---|---|---|---|
| default | Duplication | 0.987 | 0.999 |
| default | Loss | 0.812 | 6.671 |
| default | NNI | 0.920 | 3.867 |
| default | Duplication | 0.980 | 1.834 |
| default | Loss | 0.845 | 7.125 |
| default | Duplication | 0.934 | 0.609 |
| default | Loss | 0.514 | 2.386 |
| default | NNI | 0.904 | 6.078 |
| default | Duplication | 0.641 | 12.267 |
| default | Loss | -0.079 | 59.603 |
| default | Duplication | 0.966 | 0.898 |
| default | Loss | 0.670 | 1.890 |
| default | NNI | 0.910 | 2.842 |
| default | Duplication | 0.917 | 5.591 |
| default | Loss | 0.578 | 21.145 |
| default | Duplication | 0.957 | 0.847 |
| default | Loss | 0.704 | 1.442 |
| default | NNI | 0.943 | 2.709 |
| default | Duplication | 0.888 | 7.266 |
| default | Loss | 0.471 | 28.630 |
| default | Duplication | 0.963 | 0.733 |
| default | Loss | 0.728 | 1.333 |
| default | NNI | 0.955 | 2.567 |
| default | Duplication | 0.883 | 7.313 |
| default | Loss | 0.508 | 30.899 |
| default | Duplication | 0.972 | 0.846 |
| default | Loss | 0.690 | 2.184 |
| default | NNI | 0.922 | 2.547 |
| default | Duplication | 0.929 | 4.695 |
| default | Loss | 0.575 | 18.083 |
| default | Duplication | 0.948 | 0.653 |
| default | Loss | 0.644 | 1.360 |
| default | NNI | 0.941 | 2.308 |
| default | Duplication | 0.803 | 8.850 |
| default | Loss | 0.179 | 35.523 |
| default | Duplication | 0.986 | 1.039 |
| default | Loss | 0.753 | 6.826 |
| default | NNI | 0.878 | 4.544 |
| default | Duplication | 0.981 | 2.107 |
| default | Loss | 0.854 | 7.493 |
| default | Duplication | 0.987 | 0.979 |
| default | Loss | 0.804 | 6.271 |

| | | | |
|---|---|---|---|
| default | NNI | 0.908 | 3.942 |
| default | Duplication | 0.976 | 2.362 |
| default | Loss | 0.835 | 8.523 |
| default | Duplication | 0.908 | 0.623 |
| default | Loss | 0.480 | 2.971 |
| default | NNI | 0.903 | 6.699 |
| default | Duplication | 0.537 | 13.152 |
| default | Loss | -0.135 | 64.073 |
