## Supplementary material for "reconcILS: A gene tree-species tree reconciliation algorithm that allows for incomplete lineage sorting": Table S1

| Dataset | Scaled branches | Dup/Loss rate | Avg # Duplication |
|---|---|---|---|
| No error + reg branches | x1 | 0.03 | 2.844 |
| 5 errors | x1 | 0.03 | 2.844 |
| 10 errors | x1 | 0.03 | 2.844 |
| 15 errors | x1 | 0.03 | 2.844 |
| No error + long branches | x1000 | 0.00003 | 7.664 |

| Avg # Loss | Avg # NNI | reconcILS Duplication | reconcILS Loss | reconcILS NNI |
|---|---|---|---|---|
| 3.08 | 25.682 | 0.570 | 1.365 | 2.060 |
| 3.08 | 25.682 | 0.708 | 1.724 | 4.010 |
| 3.08 | 25.682 | 1.051 | 3.277 | 7.129 |
| 3.08 | 25.682 | 1.950 | 7.802 | 9.682 |
| 9.196 | 0.126 | 0.087 | 0.321 | 0.317 |

| ETE3 Duplication | ETE3 Loss |
|---|---|
| 8.510 | 33.634 |
| 10.798 | 42.520 |
| 13.408 | 54.155 |
| 15.798 | 66.273 |
| 0.052 | 0.414 |
